# Population genomics of *Bacillus anthracis* from an anthrax hyperendemic area reveals transmission processes across spatial scales and unexpected within-host diversity

**DOI:** 10.1101/2021.09.07.459365

**Authors:** Taya L. Forde, Tristan P. W. Dennis, O. Rhoda Aminu, William T. Harvey, Ayesha Hassim, Ireen Kiwelu, Matej Medvecky, Deogratius Mshanga, Henriette Van Heerden, Adeline Vogel, Ruth N. Zadoks, Blandina T. Mmbaga, Tiziana Lembo, Roman Biek

## Abstract

Genomic sequencing has revolutionized our understanding of bacterial disease epidemiology, but remains underutilized for zoonotic pathogens in remote endemic settings. Anthrax, caused by the spore-forming bacterium *Bacillus anthracis*, remains a threat to human and animal health and rural livelihoods in low- and middle-income countries. While the global genomic diversity of *B. anthracis* has been well-characterized, there is limited information on how its populations are genetically structured at the scale at which transmission occurs, critical for understanding the pathogen’s evolution and transmission dynamics. Using a uniquely rich dataset, we quantified genome-wide single nucleotide polymorphisms (SNPs) among 73 *B. anthracis* isolates derived from 33 livestock carcasses sampled over one year throughout the Ngorongoro Conservation Area, Tanzania, an area hyperendemic for anthrax. Genome-wide SNPs distinguished 22 unique *B. anthracis* genotypes within the study area. However, phylogeographic structure was lacking, as identical SNP profiles were found throughout the study area, likely the result of the long and variable periods of spore dormancy and long-distance livestock movements. Significantly, divergent genotypes were obtained from spatio-temporally linked cases and even individual carcasses. The high number of SNPs distinguishing isolates from the same host is unlikely to have arisen during infection, as supported by our simulation models. This points to an unexpectedly wide transmission bottleneck for *B. anthracis*, with an inoculum comprising multiple variants being the norm. Our work highlights that inferring transmission patterns of *B. anthracis* from genomic data will require analytical approaches that account for extended and variable environmental persistence as well as co-infection.

**Importance:** Pathogens transmitted between animals and people affect the health and livelihoods of farmers, particularly in developing countries dependent on livestock. Understanding over what distances these pathogens are transmitted and how they evolve is important to inform control strategies towards reducing disease impacts. Information on the circulation of *Bacillus anthracis*, which causes the often-lethal disease anthrax, is lacking for settings where the disease is commonplace. Consequently, we examined its genetic variability in an area in Tanzania where anthrax is widespread. We found no clear link between how closely cases were sampled and their genetic similarity. We suspect this lack of congruence is primarily driven by large-scale livestock movements, which control efforts should take into consideration. Another significant finding was the co-occurrence of multiple *B. anthracis* types within individual hosts, suggesting animals are commonly infected with a mixture of variants. This needs to be accounted for when investigating possible connections between cases.

## Introduction

Genomic data have the potential to transform our understanding of the evolution and epidemiology of pathogens of public health importance (1). However, this potential has yet to be fully harnessed for many zoonotic diseases that occur in hard-to-reach areas. Anthrax remains endemic in many low- and middle-income countries (LMICs) worldwide (2). It is a disease characterized by sudden deaths in herbivorous livestock and wildlife, and can also cause serious, potentially fatal disease in people (3). Anthrax is classified among the neglected zoonoses: a group of diseases shared by animals and people that, due to their occurrence in remote, disadvantaged communities, collectively receive less than 0.1% of international global health assistance (4). As for many neglected zoonoses, there is limited genomic data for *Bacillus anthracis*, the bacterium that causes anthrax, from endemic LMIC settings where surveillance tends to be limited. Such data could help to improve our understanding of transmission processes, such as how *B. anthracis* is spread within and between outbreaks, and ultimately contribute to more informed disease management.

The genomic diversity of *B. anthracis* has been well-described at a global scale. Isolates can be broadly divided into three major clades (A, B, C), of which A clade is the most widespread and globally dominant (5–7). Isolates from most *B. anthracis* lineages have been found across geographically widespread areas, often spanning multiple continents (7). While particular variants often predominate regionally, high lineage diversity has also been reported, including co-circulation of strains from multiple linages (8, 9). How *B. anthracis* diversity is structured at smaller scales is less well-defined. The pathogen has limited genomic diversity compared to other bacterial species (i.e. is genetically monomorphic), rendering standard genotyping methods such as multi-locus sequence typing insufficiently discriminatory (10). A hierarchical genotyping scheme known as PHRANA was therefore developed specifically for *B. anthracis*, based on quickly evolving repetitive regions nested within more phylogenetically stable markers – canSNPs – that distinguish among the major lineages (5, 11). Variants of this scheme have been used to examine the diversity of *B. anthracis* in several endemic settings globally, including in a few African countries (8, 12–14). Genome-wide SNP data would offer higher resolution for discriminating among closely-related isolates. However, whole genome sequencing (WGS) features in only a few studies of local *B. anthracis* diversity (15–18), and has rarely been conducted outside Europe (19), so the potential for phylogenomic data to be used to understand transmission patterns within hyperendemic areas has yet to be explored.

Transmission of *B. anthracis* occurs primarily through the environment. After causing the death of the animal host, vegetative bacteria are released into the environment via bodily fluids. Here, upon exposure to oxygen and cues related to a lack of nutrients, these bacteria sporulate and can persist in a dormant yet infectious state for several decades (3, 20). While the viability of spores decreases over time, new *B. anthracis* infections could theoretically arise from recent cases or cases that occurred several years or even decades previously. How this environmental persistence shapes the spatio-temporal diversity of *B. anthracis* in endemic settings has never been investigated.

In molecular epidemiological studies of bacterial pathogens, a single isolate is typically sequenced from each individual case. However, this approach fails to recognize the bacterial population diversity that may exist within the host (21, 22). Such diversity can either result from mutations that arise during infection, or from heterogeneity (multiple variants) in the inoculum, either through co-infection (exposure to multiple variants simultaneously) or superinfection (multiple exposures) (23, 24). Under those scenarios, a single isolate is unlikely to represent the overall diversity of the pathogen within the host, and as a result this approach can lead to erroneous inferences about transmission pathways (25). The importance of capturing within-host diversity is therefore increasingly recognized (26, 27). In the case of anthrax, multiple genotypes of *B. anthracis* have been previously found within individual hosts (28–30), but it remains unclear whether this represents a more widespread phenomenon.

The objective of this study was to quantify the genomic diversity of *B. anthracis* at hierarchical spatial scales within the livestock population of a hyperendemic setting. This was accomplished using a unique dataset including 1) isolates collected throughout a large (∼8,300 km^2^) area of northern Tanzania where anthrax is widespread; 2) among spatio-temporally linked cases; and 3) within individual hosts, assessing multiple isolates from the same and different sample types associated with a case (e.g. tissue, blood, soil).

## Results

### Genomic sequencing of *B. anthracis* isolates enabled assessment of diversity at hierarchical spatial scales

*Bacillus anthracis* isolates were recovered from livestock carcasses in the Ngorongoro Conservation Area (NCA), part of the Serengeti ecosystem of northern Tanzania. The 73 isolates from which WGS data were obtained were from a total of 33 carcass sites sampled throughout the NCA (Table S1). Carcasses were of the following species: sheep (n = 18), cattle (n = 7), goats (n = 4), donkey (n = 1) or unknown host (n = 3). The diversity of *B. anthracis* was assessed at multiple hierarchical spatial scales (Fig. 1). Carcasses for which detailed sampling location data were available (n = 32) could be grouped into four distinct areas within the NCA (central, north, south, east), referred to herein as ‘geographical groups’. To assess the genomic relatedness among spatio-temporally-linked cases, on four occasions, samples were collected from two carcasses either from the same or neighboring households on the same or consecutive days, which we refer to as ‘epidemiological clusters’. Three of these clusters were in the central geographical group, while the fourth was in the southern group of carcasses sampled. Finally, multiple isolates (n = 2-4) were sequenced from a single carcass for 21 carcasses. These were either i) isolates from multiple sample types (i.e. tissue, blood, swabs, soil and/or insects) from a single carcass (n = 16 carcasses) and/or ii) multiple isolates from the same sample (n = 15 carcasses). Isolates contributing to investigations of diversity at each hierarchical level are detailed in Table S1, along with individual sequence quality metrics. Mapped reads had an average depth of coverage of 85X across isolates, ranging from 24X – 245X (median 72X).

**Fig 1.**
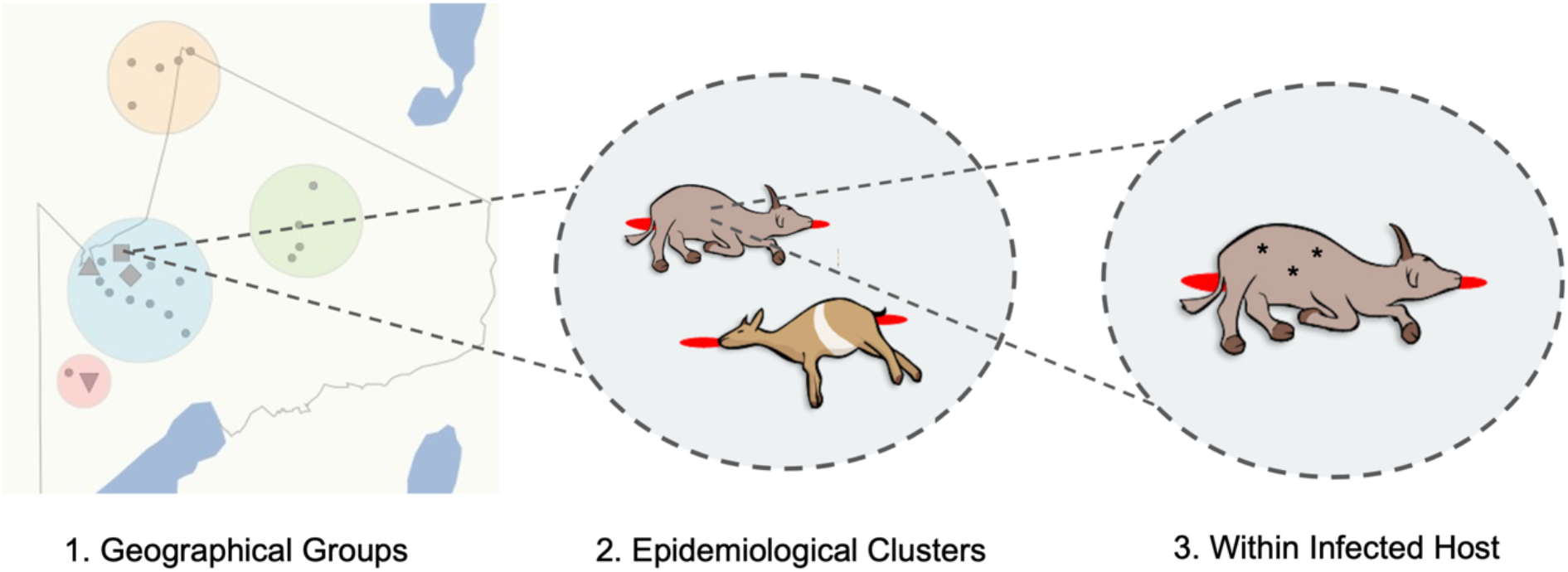
Hierarchical levels at which the genomic diversity of *Bacillus anthracis* was studied. 1. Sequenced isolates originating from livestock carcasses sampled throughout the anthrax hyperendemic Ngorongoro Conservation Area (NCA), northern Tanzania, (shown as grey dots in panel 1) were categorized into four geographical groups (colored circles); 2. A subset of these isolates were from spatio-temporally linked pairs of carcasses (n = 4, represented by grey shapes in panel 1), referred to as ‘epidemiological clusters’; 3. Multiple isolates (n = 2-4; represented by asterixis) were sequenced from individual carcasses, either originating from multiple sample types (e.g. tissue and blood), and/or multiple isolates sequenced from a single sample. The shape file for the NCA was provided by Tanzania National Parks (TANAPA) (60).

### *B. anthracis* within the NCA is limited to a single subgroup lacking clear phylogeographic signal

All *B. anthracis* isolates from the NCA were found to belong to the Ancient A subgroup of Clade A (Fig. 2). Within this subgroup, the NCA isolates formed a monophyletic clade within Cluster 3.2 as defined by Bruce et al. (7), which also contained the isolate A2075 (GenBank accession SRR2968187), isolated in 1999 from a baboon in Muhesi Game Reserve, central Tanzania. A total of 125 SNPs polymorphic among the NCA isolates and A2075 were retained for analysis, of which 13 were unique to A2075. Twenty-two unique genotypes (SNP profiles) were found among the 73 NCA isolates (Fig. 3). Based on a rarefaction analysis, the observed genotypic diversity was close to that present throughout the study area (i.e. further sampling would have been unlikely to reveal additional genotypes; Fig. S1). The maximum pairwise nucleotide difference between any two NCA-derived isolates was 49 SNPs [median = 24, interquartile range (IQR) = 10-35] (Fig. S2). There was no clear relationship between the pairwise nucleotide differences and the geographical distance between sampling locations (Fig. S3), as confirmed by a test for isolation by distance (r = 0.04, p = 0.214). Isolates from the central, eastern and southern geographical groups were observed throughout the phylogenetic tree, while all isolates from the northern sampling area were restricted to a single clade that contained the majority of NCA isolates (Fig. 3). In eight instances, identical *B. anthracis* genotypes were found in carcasses from different geographical groups, all of which involved isolates from the central geographical group and one of the other areas, with all areas implicated (Table S3). Identical or nearly identical SNP profiles (1 SNP difference) were obtained from carcasses sampled 3-5 months apart on six occasions, and 10 months apart on one occasion (Table S4).

**Fig 2.**
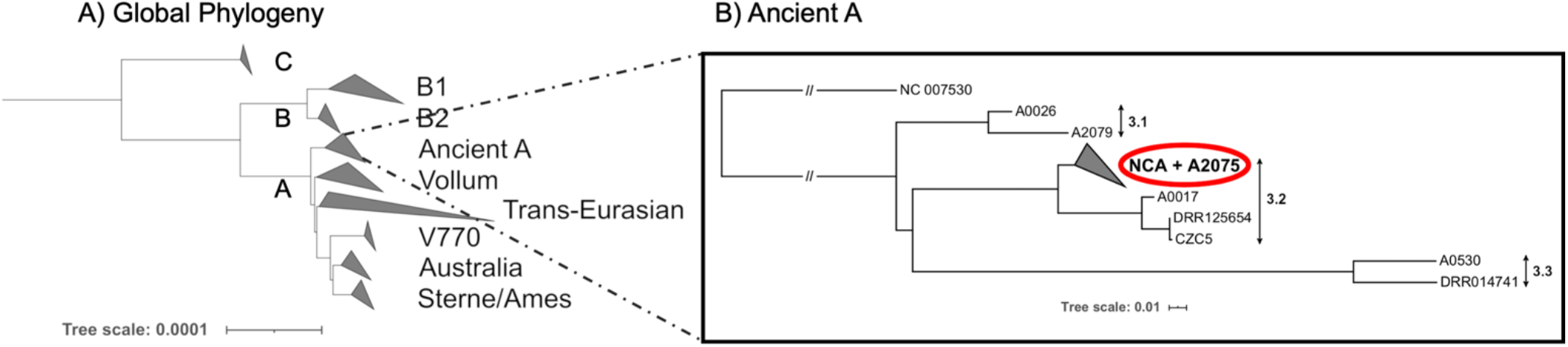
Phylogenetic position of *Bacillus anthracis* isolated from the Ngorongoro Conservation Area within the global population. A) Global phylogeny of *B. anthracis*, showing the major clades (A, B, and C) and sub-lineages. This tree was estimated based on a core single nucleotide polymorphism (SNP) phylogeny of 80 publicly available genomes (Table S2). B) Maximum likelihood phylogenetic tree of the Ancient A lineage (Cluster 3 based on Bruce et al., 2019). All isolates from the Ngorongoro Conservation Area (NCA) form a monophyletic lineage within Cluster 3.2, along with the publicly available isolate A2075 (SRR2968187) isolated in 1999 from a baboon in central Tanzania. This tree was inferred using the general time reversible (GTR) model of nucleotide substitution, using the Ames Ancestor reference genome (NC_007530) as an outgroup. Tree scales reflect number of substitutions per site.

**Fig 3.**
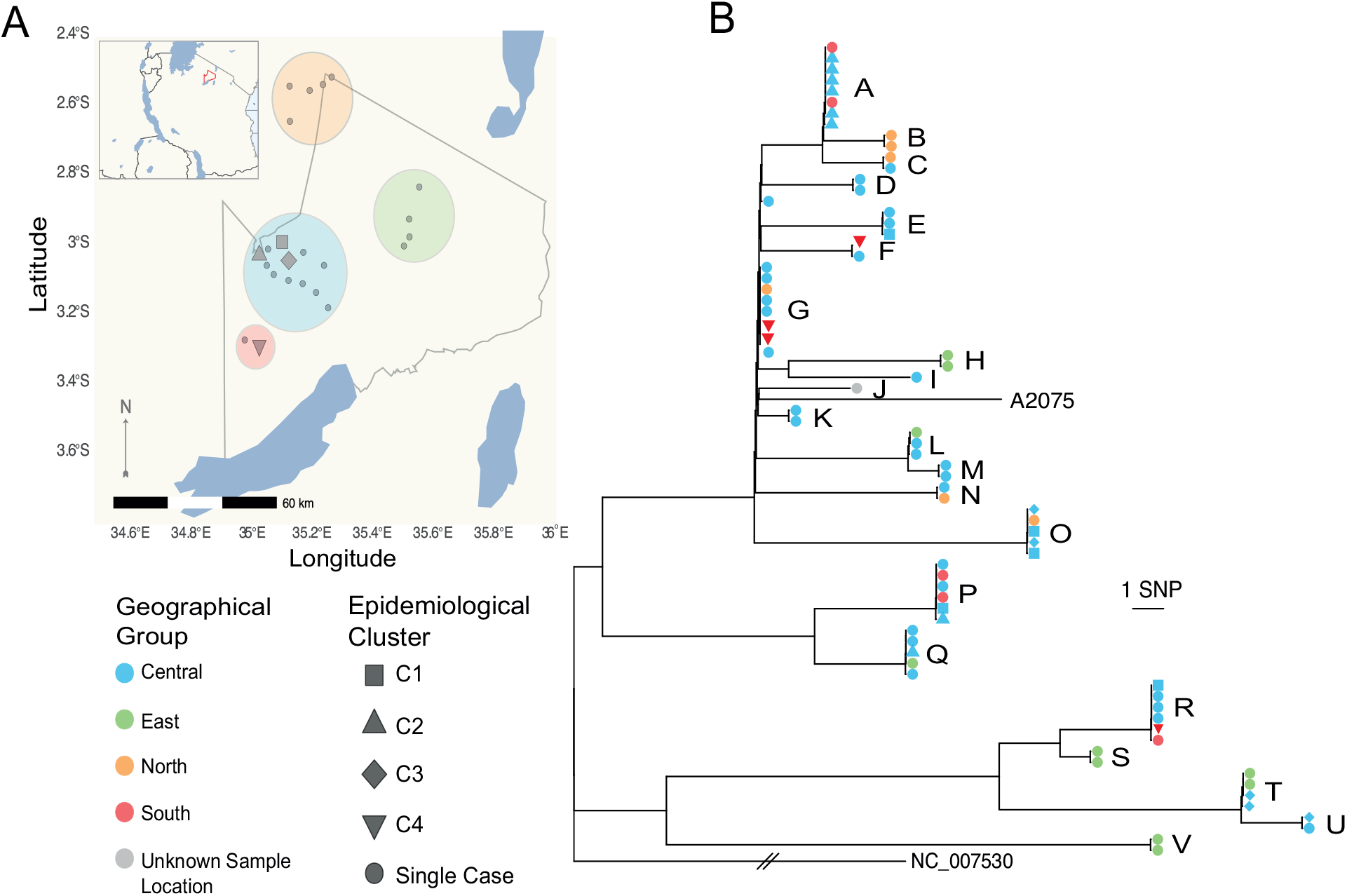
Phylogeography of *Bacillus anthracis* in the hyperendemic area of the Ngorongoro Conservation Area, Tanzania. A) Spatial distribution of carcasses from which *B. anthracis* isolates were obtained. The map outlines the Ngorongoro Conservation Area (NCA) and shows its location in northern Tanzania (outlined in red in inset). Carcasses were assigned to four geographical groups within the NCA based on spatial proximity, shown by colored circles. B) Maximum likelihood tree estimating the phylogenetic relationship among *B. anthracis* isolates from the NCA. This tree is based on an un-gapped alignment of 125 high quality core single nucleotide polymorphisms (SNPs) across the whole chromosome, rooted to the Ames Ancestor reference sequence (NC_005730) and including the publicly available isolate A2075 (SRR2968187) originating from central Tanzania. Using the more closely related isolates from Cluster 3.2 as an outgroup produced the same root position. Isolates are colored on the tree based on their collection site (geographical group) within the NCA. Epidemiological clusters of cases (pairs of carcasses sampled from the same or neighboring households on the same or consecutive days) are distinguished by symbol shape. Letters distinguish the 22 unique genotypes (SNP profiles) detected. Isolate-labelled versions of these figures can be found as Fig. S4 and S5 in the Supporting Information. The base earth, river and lake data for the map were downloaded from Natural Earth (https://www.naturalearthdata.com/). Figure was plotted in R v.3.6.1 with *ggplot2* (61), with the addition of the *sf* (62) and *ggtree* (63) packages.

### *B. anthracis* isolates from spatio-temporally linked cases are rarely genetically related

Between one and four isolates were sequenced from each carcass sampled as part of an epidemiological cluster (i.e. pair of spatio-temporally linked carcasses), resulting in between 4-8 isolates per cluster. In most cases, isolates deriving from the same epidemiological cluster were phylogenetically unrelated. Only in one of the four clusters examined (C2) did both carcasses contain isolates with identical core SNP profiles (Fig. 3B, S6 and S7). Overall, isolates from different carcasses within the same epidemiological cluster had similar numbers of SNP differences when compared with randomly selected carcasses (linked cases: median = 21, IQR = 0-33; unlinked cases: median = 23, IQR = 10-35) (Fig. 4).

**Fig 4.**
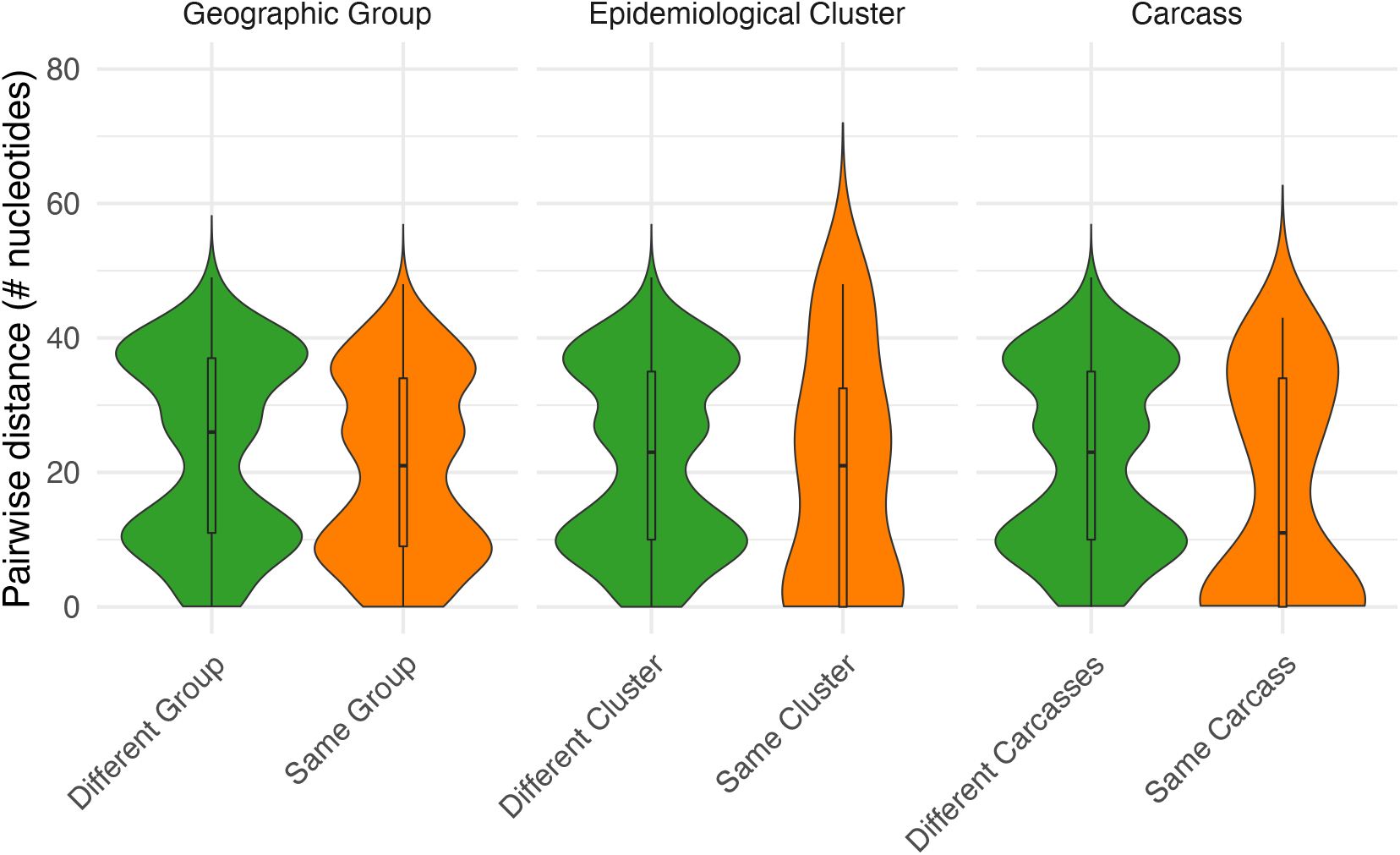
Comparative numbers of single nucleotide differences among *Bacillus anthracis* isolates from hierarchical spatial scales. Violin plots comparing pairwise nucleotide (SNP) differences between all *B. anthracis* isolates from the Ngorongoro Conservation Area versus SNP differences between isolates from i) within the same geographical group; ii) within the same epidemiological cluster (but not from the same carcass); and iii) within a single carcass. Central boxplot shows median and interquartile range, with whiskers showing minimum and maximum values.

### Within-host diversity of *B. anthracis* is similar to between-host diversity in the NCA

Overall, the number of pairwise SNP differences between isolates from the same carcass was lower than that found between isolates from different carcasses [same: median = 11, interquartile range = 0-34; different: median = 23, interquartile range = 10-35] (Fig. 4). However, isolates with multiple distinct genotypes were obtained on 15 of 21 occasions wherein multiple isolates were sequenced from the same carcass, with as many as 43 SNP differences between isolates (Fig. 5). A high level of divergence was seen, regardless of whether isolates were from the same or different sample type (e.g. multiple isolates from a tissue sample vs. isolates from tissue and soil samples, Fig. 5A). Divergent genotypes were observed in all carcasses from which three or more isolates were sequenced (12/12), and in a third (3/9) of carcasses for which two isolates were sequenced. A single SNP difference separated one isolate from two others within one carcass (AN16-83) and two SNP differences separated two isolates from a single soil sample (LNA). All other pairwise within-host SNP differences were eight or above. All of the within-host SNP profiles detected were shared with those from other carcasses.

**Fig 5.**
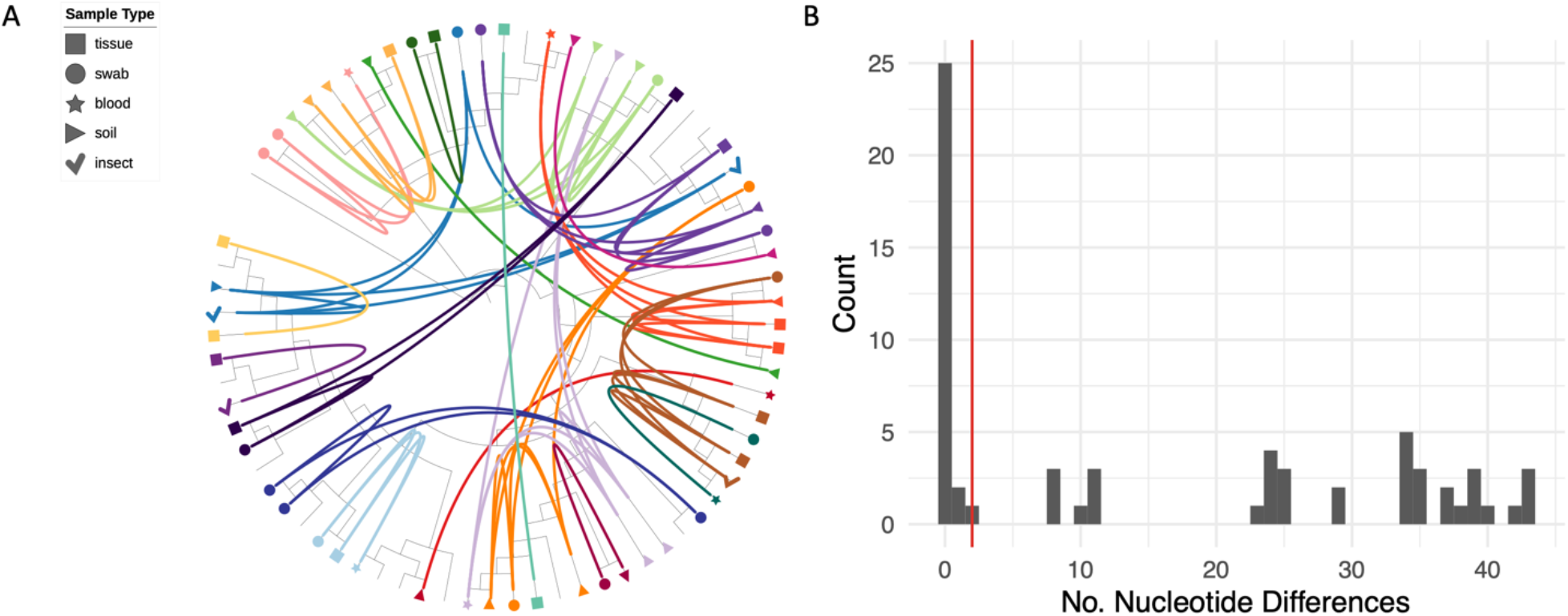
Within-host diversity of *Bacillus anthracis* among livestock in the Ngorongoro Conservation Area. A) Circularized maximum likelihood tree – based on high quality core single nucleotide polymorphisms (SNPs) – displayed as a cladogram (branch-lengths ignored), rooted to Ames Ancestor reference genome (NC_005730). Isolates from the same carcass are shown in the same color and are linked by inner connecting lines. Isolates without labels are singletons (i.e. only one isolate sequenced per carcass site). Sample type is shown by the different symbol shapes indicated in the legend. The figure was prepared using iTOL (64). For labeled taxa, see Fig. S7. B) Histogram showing the relative frequency of pairwise SNP differences among *B. anthracis* isolates collected from the same carcass. Red line shows 99% upper limit of nucleotide differences observed among sampled pairs of genomes based on simulation of within host evolution. Results suggest that almost all diversity observed within the same infected host is the result of a heterogenous inoculum.

### Observed within-host diversity is unlikely to have arisen during the course of infection

To assess the likelihood of different levels of within-host diversity arising during the course of infection, we performed simulation modelling of *B. anthracis* infection with a homogenous inoculum of varying size (1, 2, 5, 10, 20, 50 and 100 bacterial genomes), and across a range of replication cycles (up to 25 generations) using previously established mutation rates (29). With an infectious dose *d*, discrete generations and no die-off, the population size in generation *i* is therefore *d* × 2^(*i*–1)^. Averaging across simulations with varying initial doses, in the 25^th^ generation, 90.0% of the population was identical to the infecting dose, 9.53% differed by 1 SNP, and 0.489% by 2 SNPs, with higher numbers of mutations very rare (<0.01%). While stochasticity was present in early generations and particularly at low inoculum doses (Fig. S8-10), the mean proportion of genomes with various numbers of SNPs changed predictably after the initial generations (Fig. 6A-B). Extrapolating to the 40^th^ generation, then ∼84% of the population would be expected to be identical to the infecting dose, ∼15% differing by 1 SNP and ∼0.9% by 2 SNPs.

**Fig 6.**
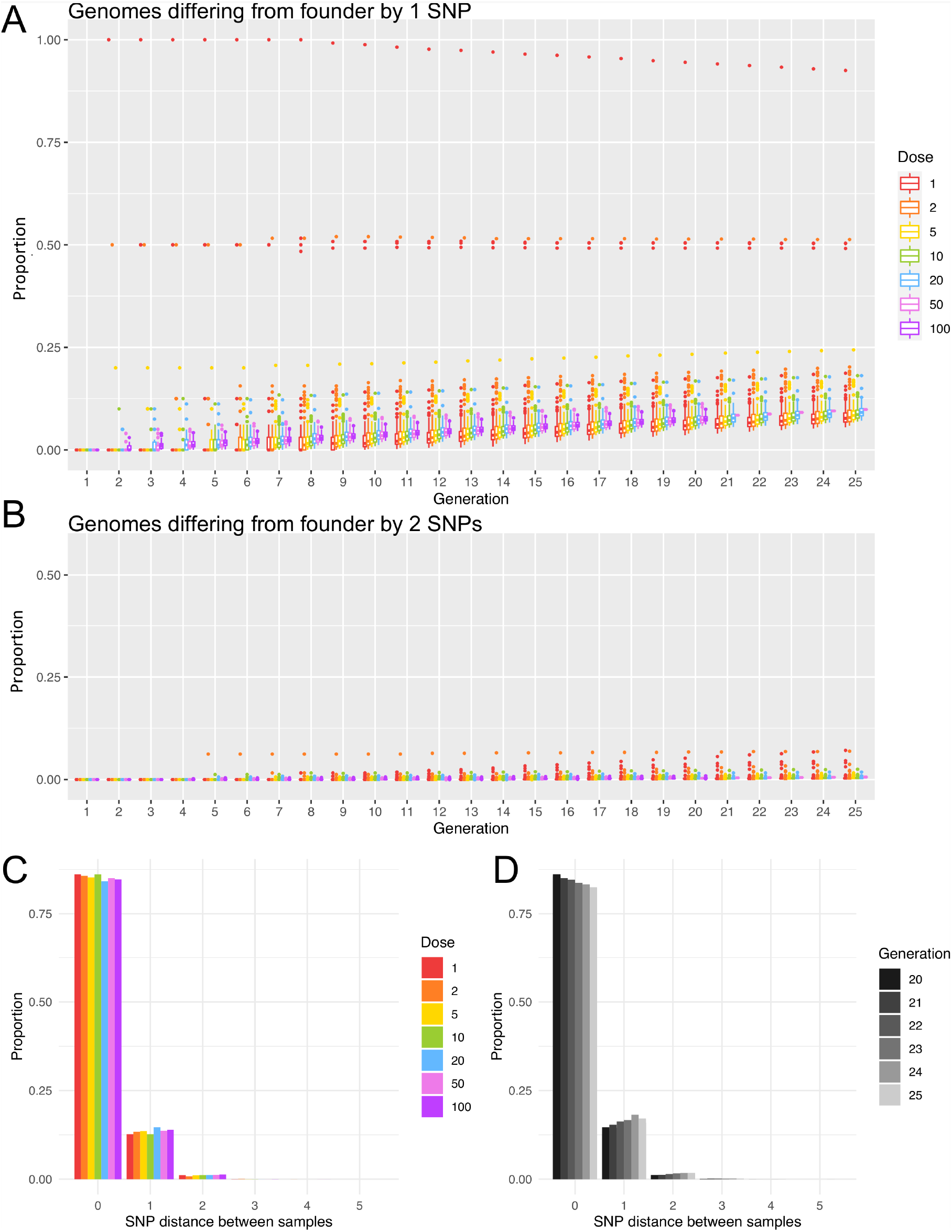
Simulation of within-host populations and sampling of resulting genetic diversity. Box and whiskers plots show the proportion of genomes across simulated populations that differ from the founding genome by either one nucleotide (A) or by two nucleotides (B). Boxes represent the interquartile range with a line showing the median and with outliers shown as points. Both boxes and outliers are colored by inoculum size (dose) according to the legend. Simulations were run for 25 generations, or 20 generations for larger inoculum sizes. Greater stochastic heterogeneity was observed for lower starting doses. Note that in (B), only outliers are visible as the proportions observed were very low. C) Bar-plot showing the relationship between inoculum size (dose) and single nucleotide polymorphism (SNP) distances between pairs of sampled genomes. Bars represent average proportions of pairwise SNP differences between genomes sampled from simulated within-host populations. Populations were simulated from various inoculum sizes and were sampled in the 20^th^ generation after 19 replication cycles. Fifty simulations were run from an initial dose of 20, and 100 simulations were run from each of the other inoculum sizes, with each simulated population sampled 100 times in each generation. D) Bar-plot showing the relationship between the number of replication cycles and SNP distances between pairs of sampled genomes. Populations were simulated for 25 generations and samples taken in generations 20-25 are represented in shades of grey according to the legend. Proportions are averaged across simulations initiated with inoculum sizes of 1, 2, 5, 10 and 20 and sampled repeatedly.

To illustrate how this within-host generated diversity would be captured in our sampling, pairs of genomes were repeatedly sampled from simulated populations and pairwise differences calculated (Fig 6C and 6D, File S1). Overall, pairwise differences greater than 2 SNPs occurred in less than 0.2% of all simulated populations (Table S5). Higher SNP distances such as those observed (Fig. 5B) are therefore unlikely to arise within-host following infection with a homogenous inoculum.

## Discussion

Genomic data for understanding the population structure and transmission patterns of bacterial zoonoses has been limited for LMIC settings where these diseases tend to have the greatest impact. Despite *B. anthracis* having limited genomic diversity in comparison with other bacterial species, WGS provided sufficient resolution to discriminate among isolates collected from within a relatively small geographic area of a few thousand square kilometers. The way in which the genomic diversity was partitioned across hierarchical spatial scales within this area has a series of novel implications for our understanding of how the pathogen is transmitted and evolves during endemic circulation.

### *B. anthracis* diversity within the hyperendemic NCA region is limited to a single Clade A sub-group

The NCA is an area where anthrax has likely been endemic for decades, if not centuries. Local community members claim that it has been an issue for their health and that of their livestock throughout living memory. Despite WGS providing sufficient discriminatory power to differentiate among individual *B. anthracis* isolates in this setting, fewer than 50 SNP differences were found among NCA isolates across the 5.2 MB chromosomal genome, highlighting the degree to which this bacterial pathogen is monomorphic. All 73 sequenced isolates formed a monophyletic cluster within the Ancient A lineage – also known as canSNP group A.Br.005/006 or A.Br.034 (6) – a subgroup of Clade A comprised mostly of isolates from south-eastern Africa. This contrasts with some previous studies of anthrax diversity in chronically endemic areas reporting co-circulation of isolates from multiple lineages using lower resolution markers (8, 9). Our study, which only included isolates collected over one year, can be considered a snapshot of recent diversity only. That being said, Tanzanian isolates genotyped in a previous study (n = 17) were also all found to belong to Ancient A clade (5), demonstrating that this particular lineage is dominant and well-established in this country. The most closely related publicly available isolate (A2075), sampled in central Tanzania approximately 300km away from our study area, differed from some NCA isolates by only 13 SNPs, which illustrates that highly related *B. anthracis* isolates can be geographically widespread. Broader, longitudinal WGS studies of *B. anthracis* across different regions of Africa will be needed to assess the actual range of individual genotypes over space and time.

### No phylogeographic signal observed despite considerable SNP diversity

Despite the considerable diversity observed (i.e. 22 unique genotypes within the sampled population), a phylogeographic signal was not detected at the scale of this study area. The finding of identical SNP profiles across distances of tens of kilometers likely reflects the ecology of anthrax in general and in our study system. First, there are few opportunities for genetic diversity to arise within the *B. anthracis* lifecycle, which is characterized by long periods of environmental dormancy in spore form, punctuated by brief interludes of a few days where it develops into its vegetative state and replicates within an infected host. It is estimated that *B. anthracis* undergoes only 20-40 replications per infection (11). Given its low mutation rate (5.2 – 8.3 × 10^−10^ mutations/site/generation) (5, 29, 31), novel mutations would likely arise in only a small proportion of *B. anthracis* infections, as also supported by our simulations. The short time of active replication within a host also means there is minimal opportunity for horizontal gene transfer with other bacteria (11), which further restricts the ability of *B. anthracis* to diversify. It is believed that *B. anthracis* rarely multiplies outside of a host, although there is some evidence for limited environmental replication (32, 33). Viability of *B. anthracis* decays exponentially over time, although infectious spores remain detectable at carcass sites at least four years after the death of the animal (34), and under favorable conditions, spores can remain viable for up to several decades (20). It is therefore reasonable to expect that spores from a single anthrax carcass with few or no novel mutations could be the source of subsequent infections over highly variable time periods, a phenomenon referred to by Sahl et al. (2016) as a ‘time capsule’. This situation violates typical molecular clock assumptions (35), wherein SNPs would be expected to arise at a relatively steady rate within the pathogen population over time. The observation that contemporary isolates from the NCA are phylogenetically basal to isolate A2075, collected nearly two decades earlier, highlights this issue. In order for molecular clock models to be applied to environmentally-persistent pathogens such as *B. anthracis* – for instance to estimate how long particular lineages have been in circulation – current analytical frameworks will need to be extended.

In addition to long and variable environmental persistence, animal movements and spatial admixture likely contribute to the lack of phylogeographic structure of *B. anthracis* within this study area. Potential sources of infection are quite spatially restricted, as the highest concentrations of viable spores are found within only a few meters of anthrax carcasses (36), in what have been termed ‘localized infectious zones’ (37). However, given that the incubation period of anthrax in livestock typically ranges from 1-14 days and potentially longer (3), extensive movement can occur between the time of infection and the animal’s death, meaning the site of sampling likely does not reflect the site of infection. In parts of rural Africa where pastoralism is the main form of agriculture, livestock are moved for various reasons, including to access water, pastures and minerals. Such daily and/or seasonal livestock movements could contribute to the observation of identical SNP profiles over distances of tens of kilometers in our study area and the lack of relationship between genetic distance and geographic distance between sampling locations. While livestock appear to be the primary drivers of *B. anthracis* transmission in the NCA, movement of infected wildlife and scavengers acting as carriers of *B. anthracis* spores could represent an additional mechanism contributing to our observations (38, 39). Further comparative genomic studies across wider areas will be essential for elucidating the geographic scales at which transmission occurs. This would help to delineate areas across which coordinated livestock vaccination campaigns should occur to avoid regular re-incursions through animal movement and trade.

### High within-host diversity is the result of simultaneous infection with multiple variants, not within-host evolution

We observed high *B. anthracis* diversity within individual hosts, essentially indistinguishable from levels of diversity found throughout the study area. Smaller numbers of SNPs (1-2) could potentially have arisen during culture; however, based on previous passaging experiments with *B. anthracis* (29), we do not expect this to have contributed in our case due to the limited number of passages performed. Alternatively, small numbers of SNPs could have arisen during the course of infection within the host. The simulations we performed suggest that isolates with >2 SNP differences between them are unlikely to be the product of within-host evolution. The regular occurrence of such differences or greater among isolates from the same carcass indicates that animals are commonly infected with a heterogenous infectious dose (i.e. ingestion of a mixture of *B. anthracis* genotypes from single or possibly multiple grazing or watering points; Fig. 7). Multiple SNP profiles were present among isolates from the same individual soil samples, supporting the occurrence of heterogeneity in a single infection source. It is also noteworthy that all within-host SNP profiles observed in this study were shared with isolates from at least one other sampled carcass (i.e. none were unique); this strongly suggests that most of the observed within-host diversity is the result of various genotypes having been present in the inoculum rather than having been generated *de novo*. This points to a wide transmission bottleneck, since a small inoculum comprising only a few spores would limit the possible diversity that could be transmitted.

**Fig 7.**
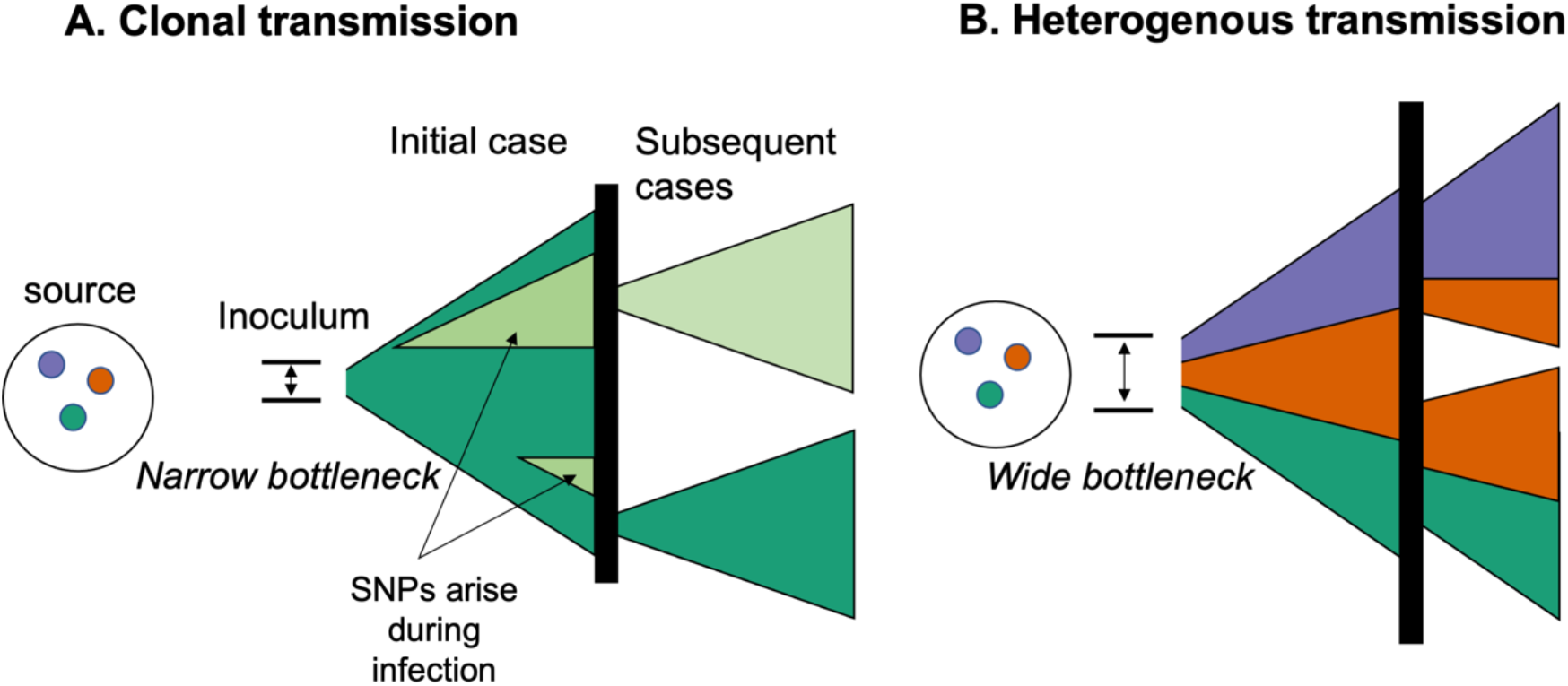
Conceptual framework for the acquisition of within-host diversity of *Bacillus anthracis*. Colors represent different genotypes of *B. anthracis* (with multiple nucleotide differences), while different shades (i.e. light green) represent single nucleotide variants arising during the course of infection. Vertical black lines represent the environment (e.g. soil, water) from which subsequent cases of anthrax arise through exposure to *B. anthracis* spores. A) Example of infections stemming from small transmission bottleneck (inoculum of few spores), resulting in primarily clonal transmission. Where the bottleneck is narrow, a limited number of genotypes could comprise the inoculum, regardless of the number of variants present in the environment. A small number of SNPs may arise during the course of infection (see Figs 5B and 6), but are unlikely to be transmitted unless they arise early during the course of infection. These variants rarely differ from the founding genotype by > 1-2 SNPs (Table S5). B) Example of infections stemming from a wide bottleneck (infectious dose with multiple spores), wherein sufficient numbers of spores comprise the inoculum such that multiple genotypes present in the environmental source may seed initial and subsequent infections. Our results strongly support heterogenous transmission (B), either from single or multiple carcass sites, and a large transmission bottleneck. Figure adapted from Ågren et al., 2014.

Our findings make an important contribution to the ongoing debate about the size of the transmission bottleneck for naturally-occurring anthrax in animals (i.e. the number of spores that give rise to a case). Recent work has proposed that founding populations may be as small as 1-3 individual spores (30). However, our findings are clearly at odds with such narrow bottlenecks, since the multiple genotypes we regularly observed within individual hosts could not have been transmitted within such a small inoculum. While our results provide no information about the exact size of the infectious dose, they align more closely with earlier suggestions of higher infectious doses, which are biologically plausible given that animals grazing at carcass sites might ingest hundreds of thousands of spores with each bite (34).

### Spatio-temporally linked anthrax cases are rarely genetically linked

Limiting phylogenetic analyses to a single isolate per host often leads to incorrect inference of transmission events (22), including the potential to overlook important epidemiological connections (40). This issue is exemplified in the current study by the spatio-temporally linked pair of cases in Cluster 2: whereas several of the isolates from both carcasses (n = 3 each) had identical SNP profiles, supporting a transmission link, both carcasses also harbored non-identical genotypes (Fig. S5 and S6). Under these circumstances, transmission links may be missed, even with complete sampling. As noted by Ågren et al. (2014), in the case of multi-clonal infections, subsequent cases may stem from different genotypes from within the founding population, masking the fact that these cases stemmed from a common source, regardless of the number of isolates sequenced among the subsequent cases. This could be the case for the other three spatio-temporally linked pairs of anthrax carcasses investigated in this study (Fig. S6), in which we only detected isolates with distinct SNP profiles. Alternatively, cases without a genetic link could be temporally linked for reasons other than exposure to a common source. For instance, animals might be more susceptible or at greater risk of exposure to infection at particular times of year, e.g. due to lower immune function related to their nutritional status and/or associated with weather extremes including prolonged rains or droughts (8). More extensive sampling of within-carcass diversity would be necessary to investigate these hypotheses and to determine whether co-occurrence of genotypes could be used to track transmission patterns.

## Conclusions

In this study, the genomic diversity of *B. anthracis* was quantified at various spatial scales within a hyperendemic setting. While WGS could discriminate among isolates within a relatively small geographic area, there was a lack of phylogeographic signal and limited genetic relatedness was observed among isolates from spatio-temporally linked cases. We hypothesize that this lack of spatial structure reflects the long-term persistence of *B. anthracis* spores in the environment, combined with extensive livestock movements related to local pastoralist practices. Based on simulations, the high within-host heterogeneity we observed points to an inoculum comprised of diverse genotypes, suggestive of a wide transmission bottleneck. Our work paves the way for studying *B. anthracis* genomic diversity and evolution within anthrax-endemic areas more broadly and to confirm the temporal and spatial scales over which genomic data are most informative for inferring transmission dynamics.

## Methods

### Study area

This study was conducted in the NCA of northern Tanzania, which covers 8,292 km^2^. Located to the south-east of Serengeti National Park, this multiple-land use area is inhabited by roughly 87,000 people (41) and one million livestock (sheep, goats and cattle) (Veterinary Officer for Ngorongoro District, pers comm). Northern Tanzania remains hyperendemic for anthrax (42), and prior to this study the NCA was recognized as a potential hotspot for this disease (43).

### Research and ethical approval

This study received ethical approval from the Kilimanjaro Christian Medical University College Ethics Review Committee (certificate No. 2050); the National Institute for Medical Research, Tanzania (NIMR/HQ/R.8a/Vol. IX/2660); Tanzanian Commission for Science and Technology (2016-95-NA-2016-45); and the College of Medical Veterinary and Life Sciences ethics committee at the University of Glasgow (200150152). It also received permission under Section 20 of the Animal Diseases Act 35 (1984) at the University of Pretoria, South Africa (Ref 12/11/1/1/6).

### Sample collection

Samples were collected between May 2016 and April 2017 inclusive through active surveillance by a dedicated field team. Sudden deaths in animals reported by community members throughout the NCA were investigated and samples were collected when anthrax was suspected (File S1). When available, the following samples were collected: a piece of skin tissue (tip of the ear if the carcass was still intact, or a piece of hide if the carcass had already been opened); whole blood; swab of blood or body fluid at natural orifices; blood- or body fluid-soaked soil from below the carcass; and insects found on or around the carcass. Various metadata were recorded, including the species of animal affected and the location of sampling (Table S1, File S1). All samples were stored at ambient temperature for up to six months at local veterinary facilities until transport to the Kilimanjaro Clinical Research Institute (KCRI) in Moshi, Tanzania for molecular diagnostics, as previously described (44), with aliquots shipped to the University of Pretoria, South Africa, for selective culture and DNA extraction from *B. anthracis* isolates.

### Selective culture, DNA extraction and sequencing

Sample pretreatment (i.e. to inhibit competition from heat-sensitive bacteria) is described in File S1. Sample homogenates (100 µL) were plated onto both polymyxin-EDTA thallous acetate (PET) selective media and 5% sheep blood agar (SBA). These were incubated at 37 °C overnight and the plates inspected for growth after 15 – 24 hours incubation. The PET was then further incubated and inspected at 48 hours. Suspect *B. anthracis* colonies based on typical morphological characteristics were sub-cultured onto SBA for purification and identification (File S1). In parallel, a single colony was streaked onto a new purity plate for nucleic acid extraction. In some instances multiple isolates were selected from the same sample where the colonies demonstrated differences in morphology but were identified on the same plate and met the selection criteria (File S1).

DNA extracts from 75 isolates from 33 carcasses were submitted for library preparation and sequencing at MicrobesNG (Birmingham, UK). Libraries were prepared using the Nextera XT v2 kit (Illumina, San Diego, USA) and sequenced on the Illumina HiSeq platform, generating 250 base pair paired-end reads.

### Bioinformatics and genomic analyses

Reads were adapter trimmed by MicrobesNG using Trimmomatic v0.30 (45), and basic statistics determined using QUAST (46) (Table S1). Bacterial species identification was confirmed using Kraken (47). Based on these quality metrics, sequences from two isolates were excluded from further analyses: one due to a low number of reads (<40,000), and another that was identified as *B. cereus*. There were some further indications that not all cultures were pure *B. anthracis*, despite multiple rounds of sub-culture (File S1, Table S1). A reference-based mapping approach and strict variant filtering criteria were implemented to minimize the issues associated with the sequence quality while making use of as much of the data as possible.

Read mapping and variant calling were performed on the CLIMB computing platform for microbial genomics (48). Trimmed reads were aligned to the chromosome of the Ames Ancestor reference genome (NC_007530) using bwa-mem (version 0.7.17). Picard was used to mark and remove duplicate reads, add read group information, and index the bam files (49). Quality metrics for read mapping were obtained using Qualimap (50) (Table S1). SNPs were detected in individual isolates by VarScan v2.4.4 (51) with parameters set as follows: minimum read depth of 4; minimum base quality of 20; variant allele frequency ≥ 0.95. Subsequent SNP curation steps are described in File S1. Custom python scripts for the assessment of read mapping SNP metrics data, variant site filtering and generation of variant call and alignment files (source codes with description of their functionality and usage), along with the final variant call and multiple sequence alignment files are available on GitHub (https://github.com/matejmedvecky/anthraxdiversityscripts).

The alignment of concatenated SNPs was analyzed using ModelFinder to determine the most appropriate model of nucleotide substitution (52). Subsequently, a maximum-likelihood phylogeny was estimated in IQ-Tree (53) under the Kimura-3-parameters (K3P) model, using 1000 ultrafast bootstrap replicates (54). A distance matrix detailing SNP differences between isolates was constructed using snp-dists v0.6 (https://github.com/tseemann/snp-dists). The distance between GPS points was calculated using the pointDistance command in the R package *raster* (55). Isolation by distance was tested using Mantel test to assess the correlation between SNP distance and Euclidean geographic distance within the R package *adegenet* (56). All program versions and commands used, the distance matrix, as well as small custom scripts are available on GitHub (https://github.com/tristanpwdennis/anthrax_diversity).

To place the newly sequenced isolates within the global phylogeny of *B. anthracis*, 80 WGS from GenBank were accessed (Table S2), and a core genome alignment generated using Parsnp v1.1.2 (57). The resulting phylogeny indicated that NCA isolates belong to the Ancient A lineage. To further resolve the diversity among NCA isolates compared with other publicly available isolates from the same lineage, reads from eight additional isolates available on SRA from a study by Bruce et al. 2019 were accessed using fastq-dump from the SRA-toolkit: all isolates (n=4) belonging to the 3.2 linage, and two arbitrarily selected isolates from each the 3.1 and 3.3 lineages (Table S2). These were run through our SNP-calling pipeline as described above, resulting in a sequence alignment file free of -/N characters, which was used to infer a phylogeny in RAxML v8.2.11 (58) using a GTR model of nucleotide substitution, and using the Ames Ancestor reference genome as an outgroup.

### Simulation modelling

The initial number of genomes in the inoculum was varied between one and 100 genomes to reflect uncertainty in anthrax infectious dose. In each bacterial generation, genomes underwent a round of replication followed by cell division and accordingly population size doubled in each generation. The number of mutations occurring in the replication of each genome was drawn from a Poisson distribution parameterized to reflect the estimated *B. anthracis* mutation rate (i.e. λ = 0.004316). This genome-level mutation rate is based on the genome size of 5.2 million base pairs and a mutation rate of 8.3 × 10^−10^ mutations per site (29); this represents an upper estimate of the mutation rate and was chosen as we were interested in estimating the upper limits on reasonable expectation of diversity emerging during the course of an infection initiated by a homogenous dose. Simulations were carried out for up to 30 generations (replication cycles), and results extrapolated to 40 generations, proposed to be the upper limit on the number of replication cycles during an infection (11). Details of simulations run are provided in File S1. Pairs of genomes were repeatedly sampled from these simulated populations and the count of mutations separating each pair was calculated. Sampling was performed 100 times per generation for each simulation. Simulations, sampling of simulated populations, and linear models summarizing trends in the outcome of these processes were performed in R (59).

## Data availability

Raw sequencing reads are available on European Nucleotide Archive SRA under accession number PRJEB45684.

## Acknowledgements

We are grateful for all the support received for this research, particularly from the NCA community and authorities. We thank the Ngorongoro District Council, Ngorongoro Conservation Area Authority, District Veterinary Office, Tanzania Wildlife Research Institute and members of our field team – Sabore Ole Moko, Sironga Nanjicho, Kadogo Lerimba and Godwin Mshumba – for assistance with this study. We also thank the Directorate of Veterinary Services, Ministry of Agriculture, Livestock and Fisheries, and Ministry of Health, Community Development, Gender, Elderly and Children for their support. Finally, we thank Yi Xuan Chew and Nichith Kollanandi Ratheesh for initial assessments of the publicly available genomic data. Establishment and maintenance of the Zoonoses laboratory at KCRI (BTM) was supported by the BBSRC (BB/J010367/1), BBSRC Zoonoses and Emerging Livestock Systems (BB/L017679, BB/L018926, BB/L018845), and the Wellcome Trust (096400/Z/11/Z).

## Supporting Information

**Fig S1. Rarefaction curve of *Bacillus anthracis* genotypic diversity within the study area**. Based on inclusion of all isolates sequenced (n = 73, left) and on all isolates that were not sampled as part of an epidemiological cluster (n = 51, right), which might be expected to be non-independent. Results suggest genotypes within this population have been exhaustively sampled (i.e. that further sampling would not be expected to reveal additional genotypes). Figure generated in R package iNEXT (65).

**Fig S2. Histogram showing the relative frequency of pairwise nucleotide (SNP) differences among *B. anthracis* isolates**. Includes 73 isolates from the Ngorongoro Conservation Area, northern Tanzania.

**Fig S3. Scatter plots showing the number of nucleotide differences as a function of geographic difference between the sampling locations**. Geographic distance (in meters) is shown on the x-axis versus nucleotide differences on the y-axis, with each point representing a pair of isolates. A) All *Bacillus anthracis* isolates from the study area. B) The same relationship is observed when limited to isolates from the dominant clade; this was done in order to account for deeper divergences potentially obscuring patterns.

**Fig S4. Locations of carcasses sampled within the Ngorongoro Conservation Area, northern Tanzania**. Each carcass is assigned a unique identifier (Table S1). Epidemiological clusters are the same as those explained for Fig 3 and Fig S5.

**Fig S5. Phylogenetic relationship among *Bacillus anthracis* isolates from the Ngorongoro Conservation Area**. Estimated through maximum likelihood, based on high quality core SNPs. Geographic groups and epidemiological clusters are the same as those explained for Fig 3. Each isolate is attributed to a carcass ID, with the sample type indicated following the underscore (B = blood, D = soil, I = insect, S = swab, T = tissue). A number follows the sample type if more than one isolate was sequenced from the same sample.

**Fig S6. Genotypes of *Bacillus anthracis* observed in isolates from within and between pairs of carcasses from the same epidemiological clusters (C1-4)**. Genotype letters correspond to those in Fig 3 and Fig S5. Individual carcasses are numbered /1 or /2. In cluster C3, two isolates were from a soil sample (C3/1) collected at the same household as the two cases (C3/2 and C3/3); genotype T from this soil sample was shared with an isolate from C3/2. Otherwise, only in C2 was there evidence of a shared genotype between pairs of carcasses (genotype A). Thus, the level of sampling conducted here (1-4 isolates per carcass) did not produce evidence for the same combinations of genotypes being found among linked carcasses.

**Fig S7. Within-host diversity of *Bacillus anthracis* isolated from livestock in the Ngorongoro Conservation Area of northern Tanzania**. This circularized maximum likelihood tree – based on high quality core single nucleotide polymorphisms – is displayed as a cladogram (branch-lengths ignored). Isolates from the same carcass are shown in the same colour and linked by inner connecting lines. Isolates in black are singletons (i.e. only one isolate sequenced per carcass site). The figure was prepared using ITOL (64).

**Fig S8. Proportion of simulated within-host populations identical to the inoculating genome over 25 generations**. Populations were simulated from homogenous inoculating doses of varying size (A-G) and each line tracks a single simulated population through generations. Represented are 100 simulations run for 25 generations from doses 1, 2, 5 and 10; 50 simulations run for 25 generations (dose 20); 100 simulations run for 20 generations from doses 50 and 100 and 7 simulations run for 25 generations from inoculum dose of 50.

**Fig S9. Proportion of simulated within-host populations differing from the inoculating genome by one nucleotide (SNP)**. Populations were simulated from homogenous inoculating doses of varying size (A-G) and each line tracks a single simulated population through generations. Represented are 100 simulations run for 25 generations from doses 1, 2, 5 and 10; 50 simulations run for 25 generations (dose 20); 100 simulations run for 20 generations from doses 50 and 100 and 7 simulations run for 25 generations from inoculum dose of 50.

**Fig S10. Proportion of simulated within-host populations differing from the inoculating genome by two nucleotides (SNPs)**. Populations were simulated from homogenous inoculating doses of varying size (A-G) and each line tracks a single simulated population through generations. Represented are 100 simulations run for 25 generations from doses 1, 2, 5 and 10; 50 simulations run for 25 generations (dose 20); 100 simulations run for 20 generations from doses 50 and 100 and 7 simulations run for 25 generations from inoculum dose of 50.

**Table S1. Metadata and sequence quality metrics of *Bacillus anthracis* isolates included in this study**. Each isolate is attributed to a carcass ID, with the sample type indicated following the underscore (B = blood, D = soil, I = insect, S = swab, T = tissue). A number follows the sample type if more than one isolate was sequenced from the same sample.

**Table S2. Publicly available *Bacillus anthracis* isolates included to contextualize those from the Ngorongoro Conservation Area**. Sheet 1: Publicly available global collection of assembled *B. anthracis* genomes. Sheet 2: Publicly available sequence data for Ancient A isolates, available on SRA.

**Table S3. Identical isolates found from different geographical groups**.

**Table S4. Identical or nearly identical isolates from carcasses sampled several months apart**.

**Table S5. Single nucleotide polymorphism (SNP) distances among pairs of isolates sampled from simulated within-host populations of *Bacillus anthracis***. Proportion of pairs of isolates in evolved populations with different SNP distances across varying initial inoculum size (dose), sampled in generations 20 and 25, and mean SNP differences across sampled pairs.

**File S1. Supplementary methods and results**.

