## Supplementary material for "Population genomics of *Bacillus anthracis* from an anthrax hyperendemic area reveals transmission processes across spatial scales and unexpected within-host diversity"

#### Supplementary Figures

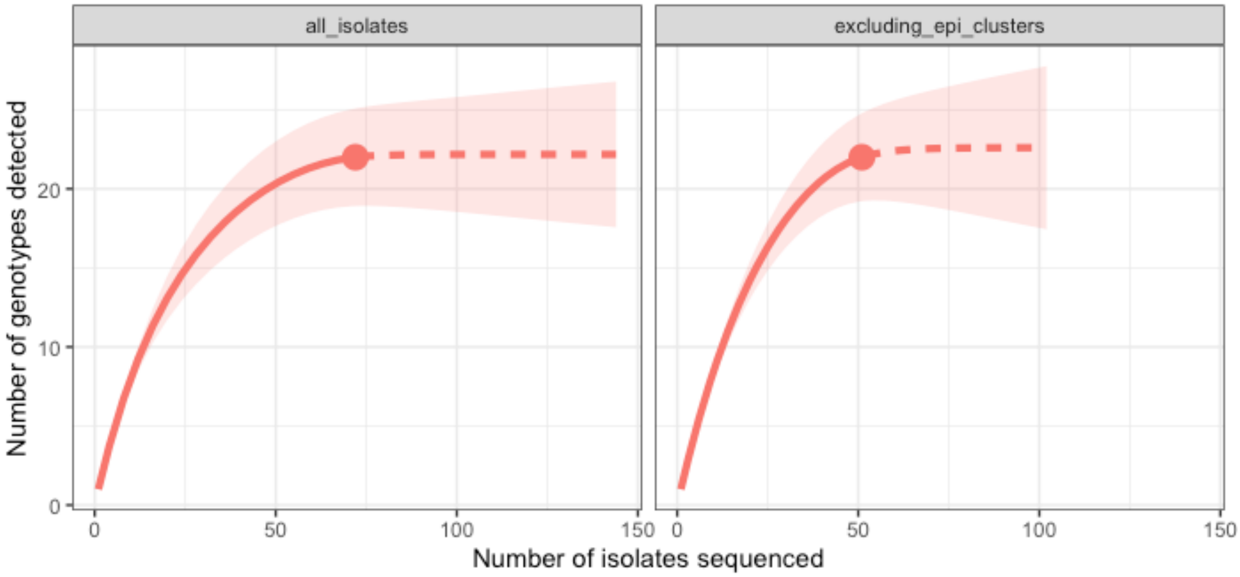


**Fig S1. Rarefaction curve of *Bacillus anthracis* genotypic diversity within the study area.** Based on inclusion of all isolates sequenced (n = 73, left) and on all isolates that were not sampled as part of an epidemiological cluster (n = 51, right), which might be expected to be non-independent. Results suggest genotypes within this population have been exhaustively sampled (i.e. that further sampling would not be expected to reveal additional genotypes). Figure generated in R package iNEXT (1).

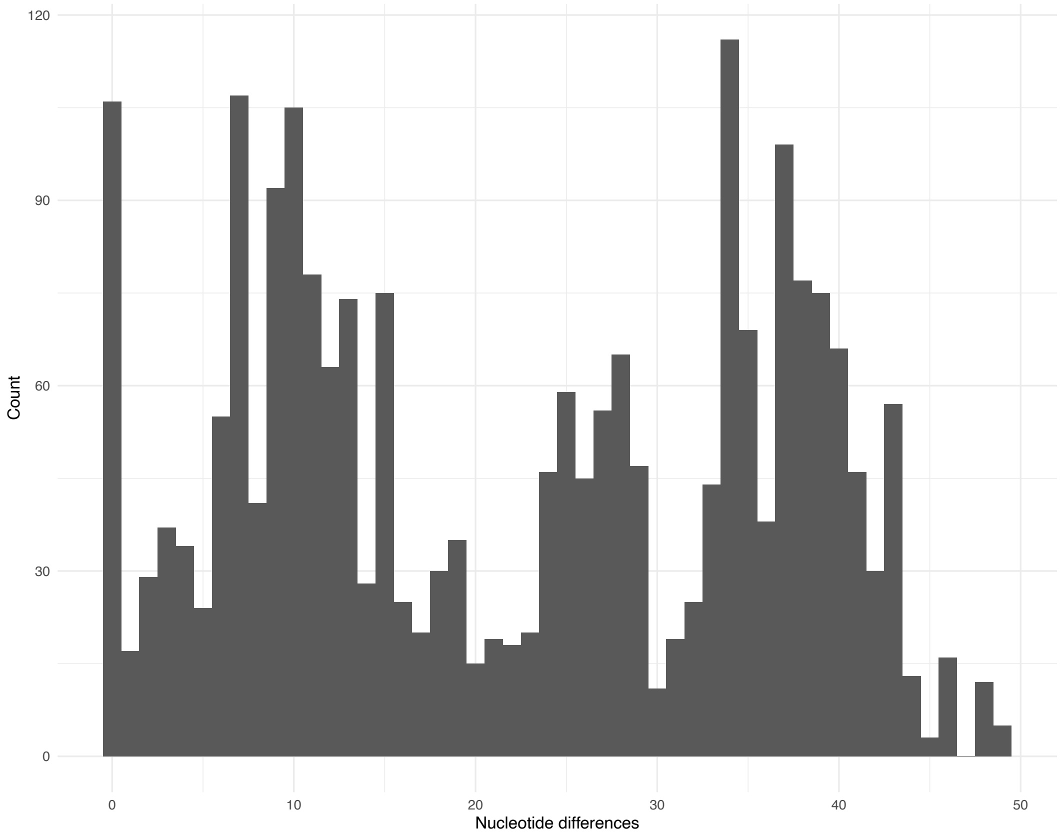


**Fig S2. Histogram showing the relative frequency of pairwise nucleotide (SNP) differences among *B. anthracis* isolates.** Includes 73 isolates from the Ngorongoro Conservation Area, northern Tanzania.

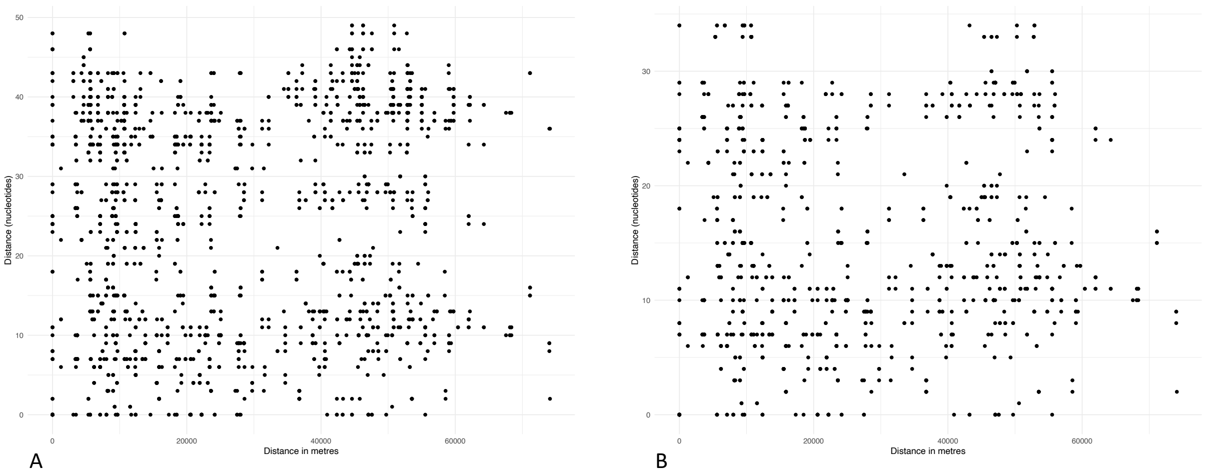


**Fig S3. Scatter plots showing the number of nucleotide differences as a function of geographic difference between the sampling locations.** Geographic distance (in meters) is shown on the x-axis versus nucleotide differences on the y-axis, with each point representing a pair of isolates. A) All *Bacillus anthracis* isolates from the study area. B) The same relationship is observed when limited to isolates from the dominant clade; this was done in order to account for deeper divergences potentially obscuring patterns.

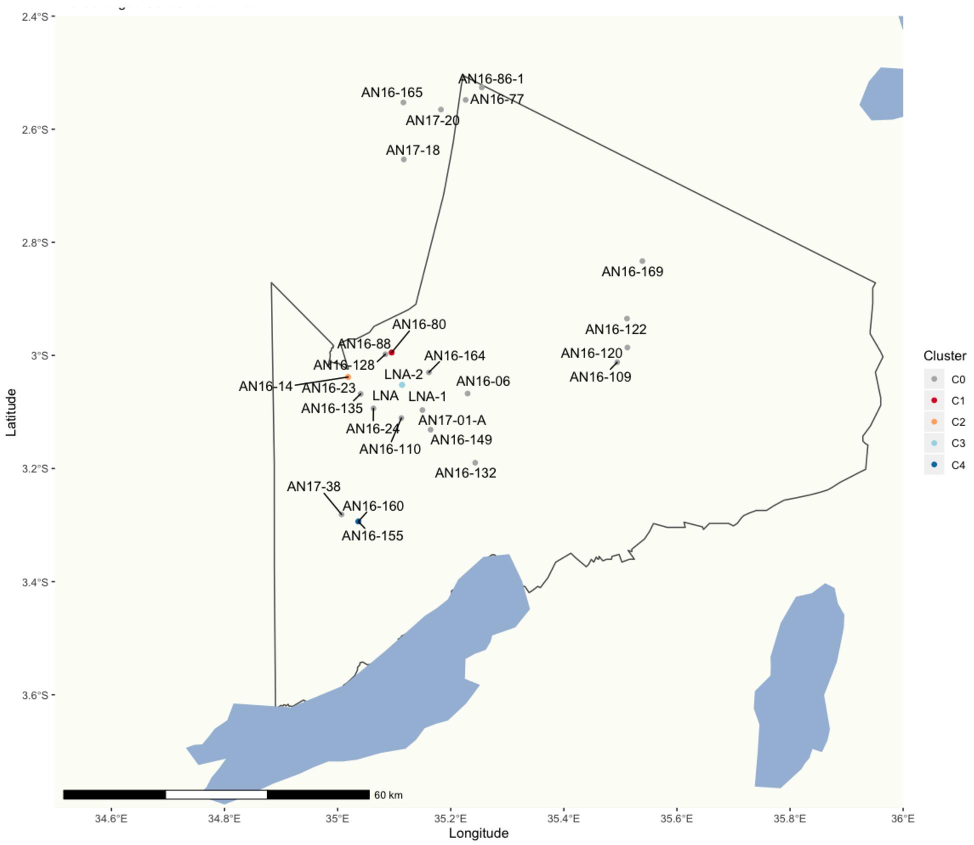


**Fig S4. Locations of carcasses sampled within the Ngorongoro Conservation Area, northern Tanzania.** Each carcass is assigned a unique identifier (Table S1). Epidemiological clusters are the same as those explained for Fig. 3 and Fig. S6.

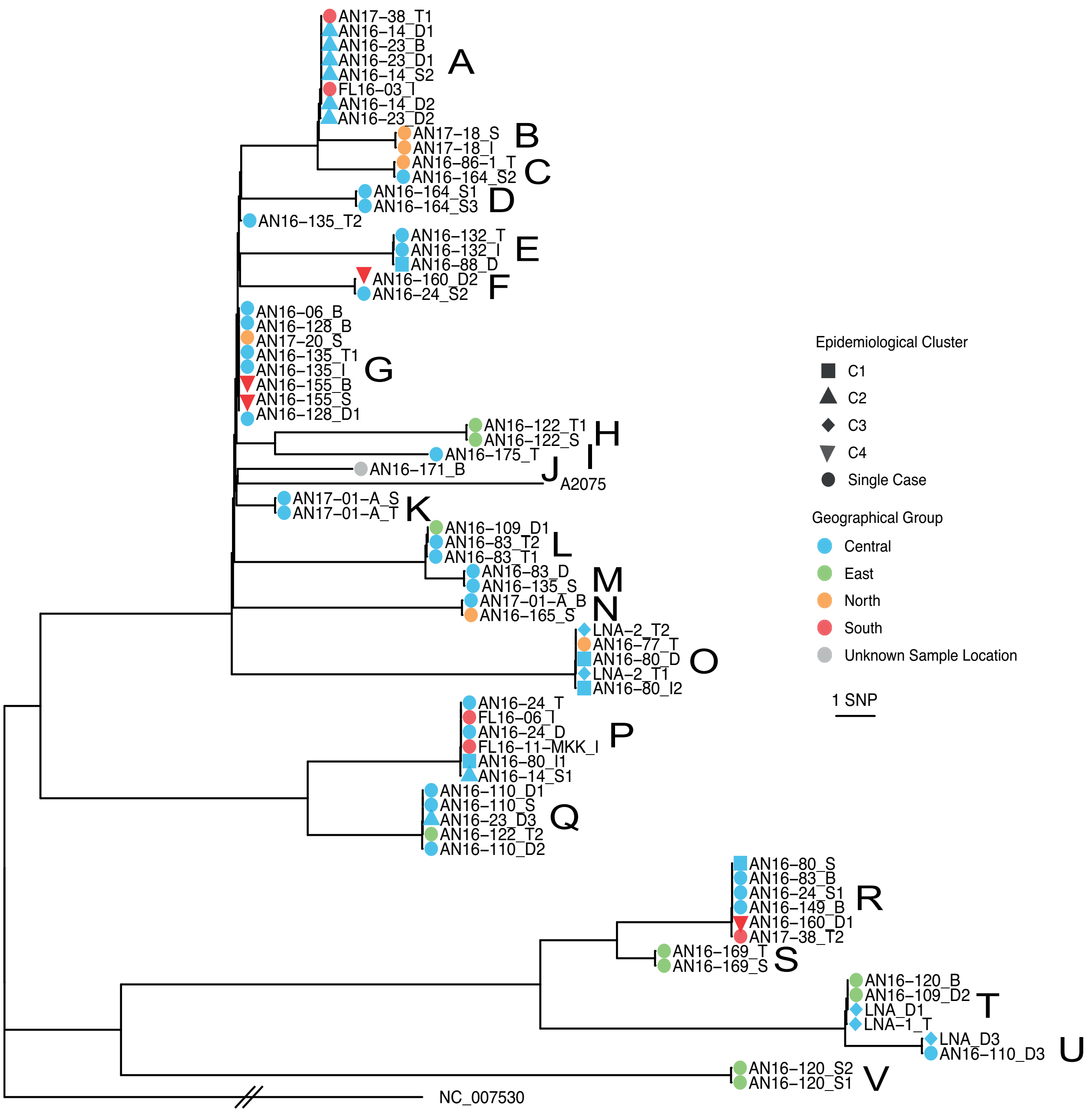


**Fig S5. Phylogenetic relationship among *Bacillus anthracis* isolates from the Ngorongoro Conservation Area.** Estimated through maximum likelihood, based on high quality core SNPs. Geographical groups and epidemiological clusters are the same as those explained for Fig 3. Each isolate is attributed to a carcass ID, with the sample type indicated following the underscore (B = blood, D = soil, I = insect, S = swab, T = tissue). A number follows the sample type if more than one isolate was sequenced from the same sample.

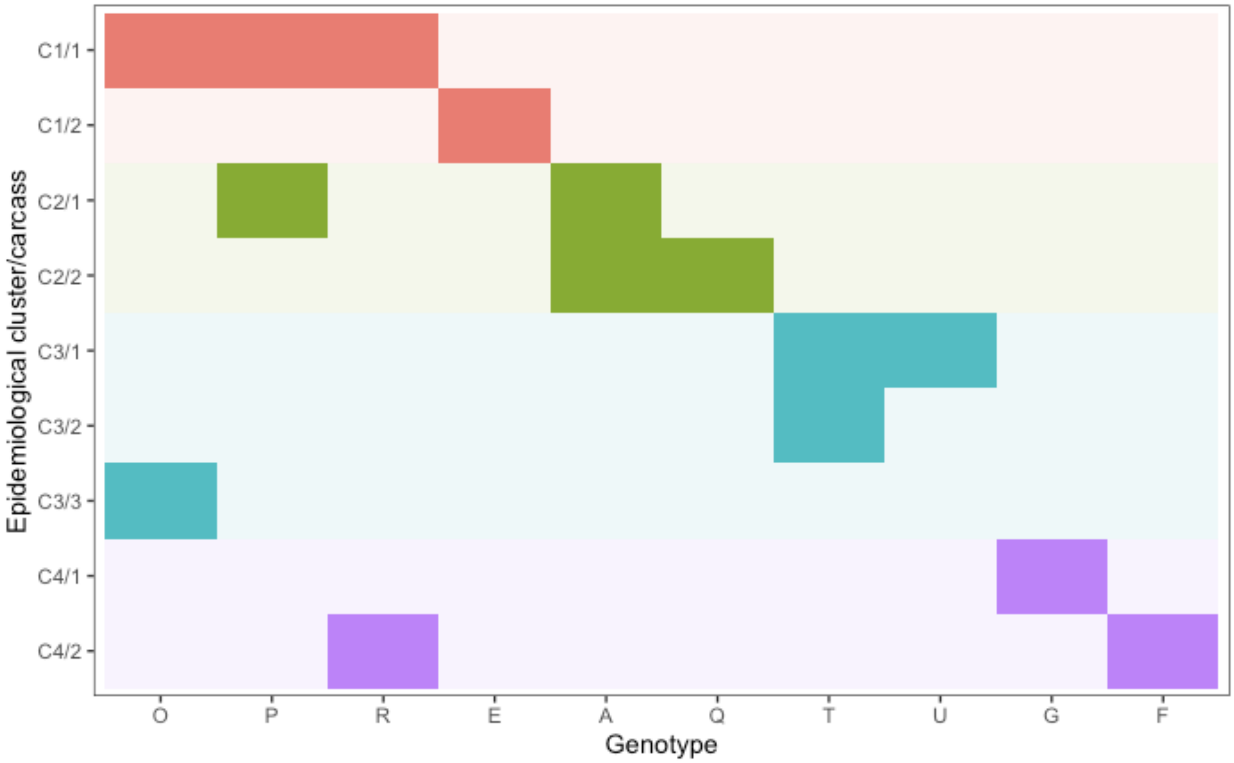


**Fig S6. Genotypes of *Bacillus anthracis* observed in isolates from within and between pairs of carcasses from the same epidemiological clusters (C1-4).** Genotype letters correspond to those in Fig. 3 and Fig. S5. Individual carcasses are numbered /1 or /2. In cluster C3, two isolates were from a soil sample (C3/1) collected at the same household as the two cases (C3/2 and C3/3); genotype T from this soil sample was shared with an isolate from C3/2. Otherwise, only in C2 was there evidence of a shared genotype between pairs of carcasses (genotype A). Thus, the level of sampling conducted here (1-4 isolates per carcass) did not produce evidence for the same combinations of genotypes being found among linked carcasses.

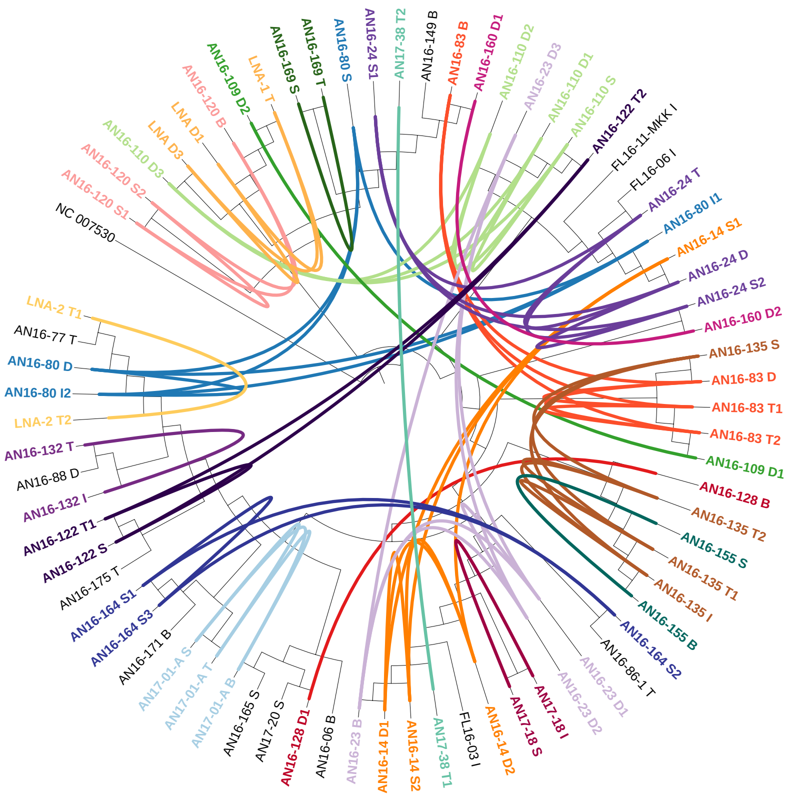


**Fig S7. Within-host diversity of *Bacillus anthracis* isolated from livestock in the Ngorongoro Conservation Area of northern Tanzania.** This circularized maximum likelihood tree – based on high quality core single nucleotide polymorphisms – is displayed as a cladogram (branch-lengths ignored). Isolates from the same carcass are shown in the same colour and linked by inner connecting lines. Isolates in black are singletons (i.e. only one isolate sequenced per carcass site). The figure was prepared using ITOL (2).

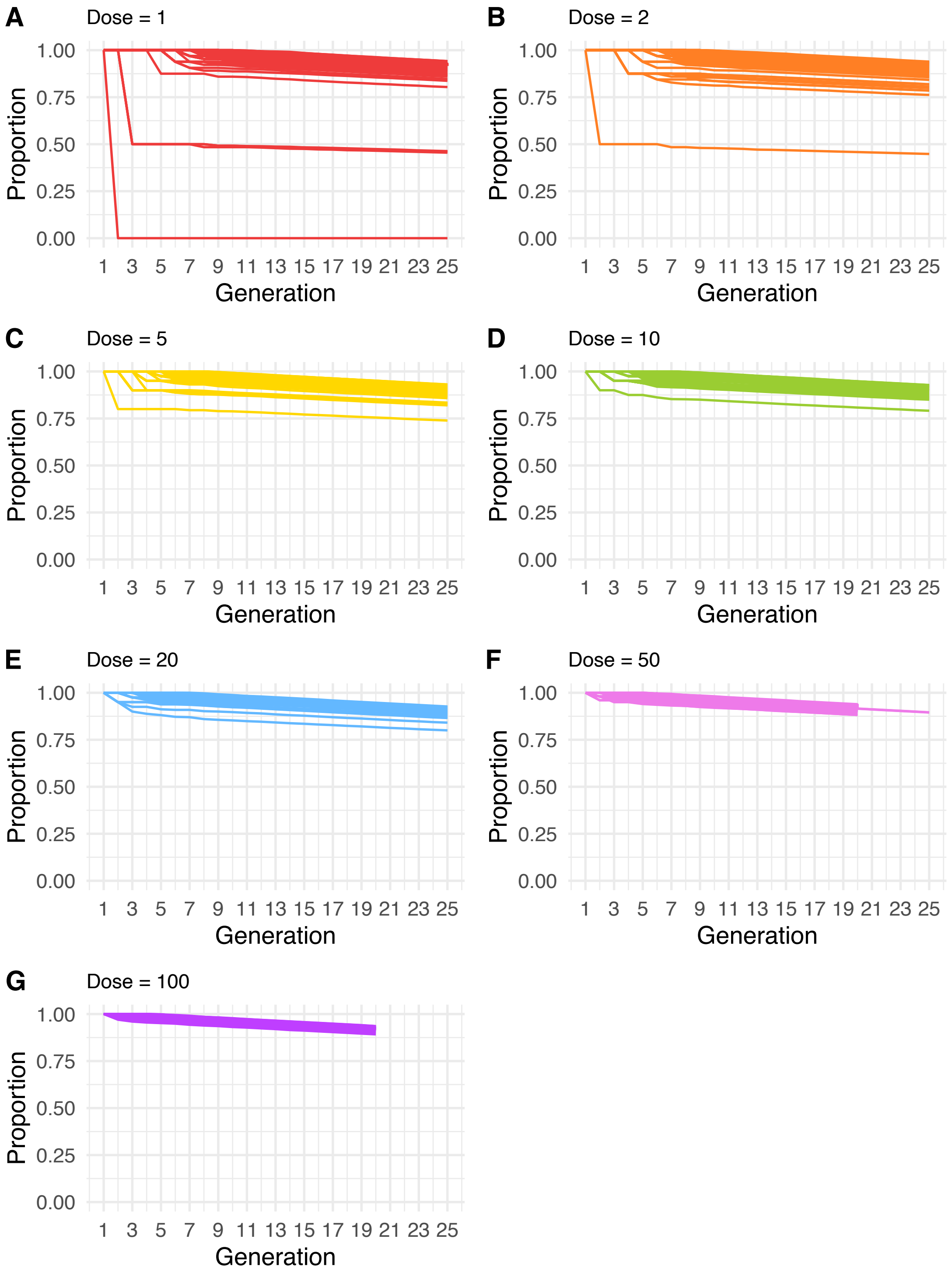


**Fig S8. Proportion of simulated within-host populations identical to the inoculating genome over 25 generations.** Populations were simulated from homogenous inoculating doses of varying size (A-G) and each line tracks a single simulated population through generations. Represented are 100 simulations run for 25 generations from doses 1, 2, 5 and 10; 50 simulations run for 25 generations (dose 20); 100 simulations run for 20 generations from doses 50 and 100 and 7 simulations run for 25 generations from inoculum dose of 50.

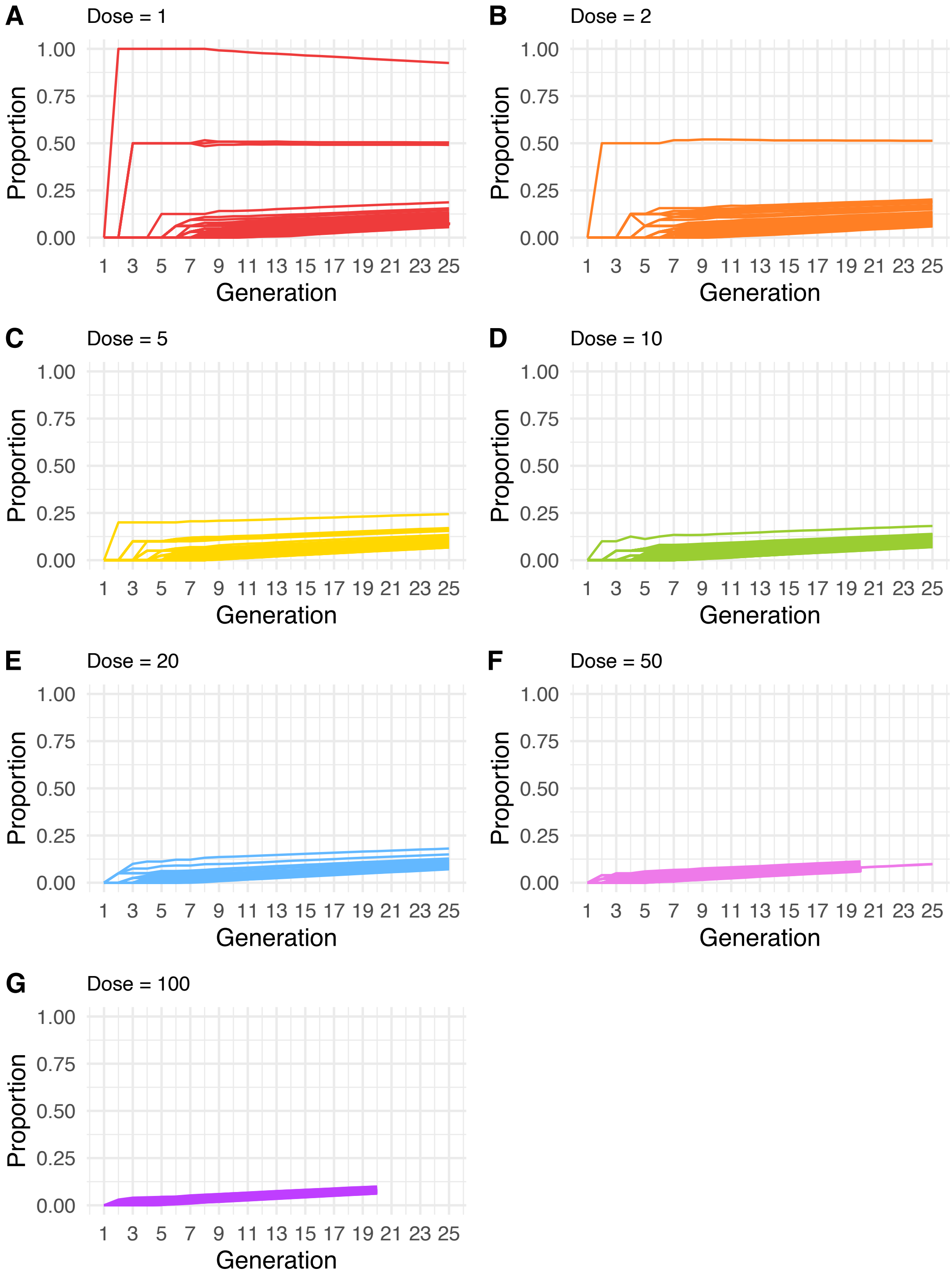

**Fig S9. Proportion of simulated within-host populations differing from the inoculating genome by one nucleotide (SNP).** Populations were simulated from homogenous inoculating doses of varying size (A-G) and each line tracks a single simulated population through generations. Represented are 100 simulations run for 25 generations from doses 1, 2, 5 and 10; 50 simulations run for 25 generations (dose 20); 100 simulations run for 20 generations from doses 50 and 100 and 7 simulations run for 25 generations from inoculum dose of 50.

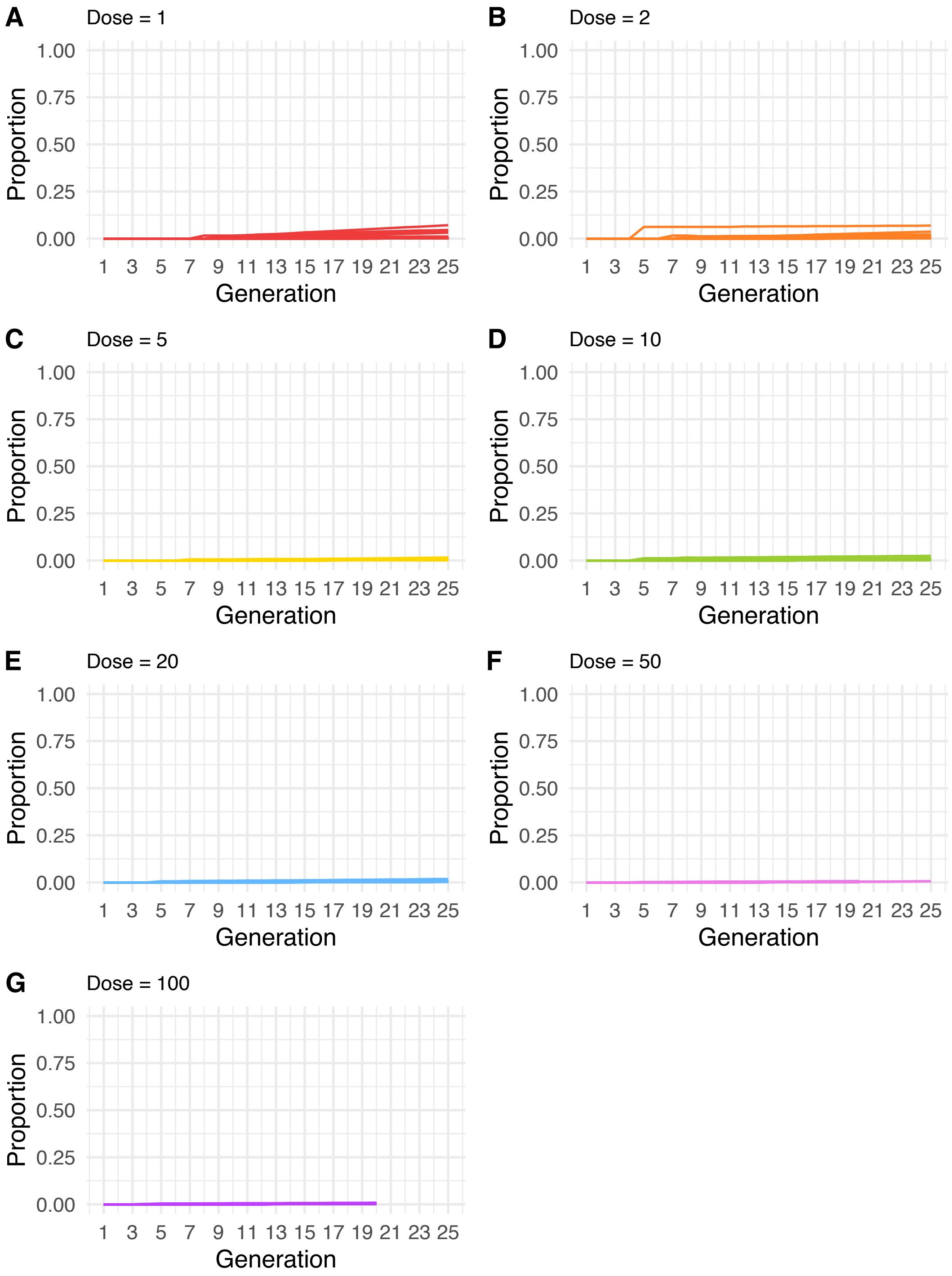


**Fig S10. Proportion of simulated within-host populations differing from the inoculating genome by two nucleotides (SNPs).** Populations were simulated from homogenous inoculating doses of varying size (A-G) and each line tracks a single simulated population through generations. Represented are 100 simulations run for 25 generations from doses 1, 2, 5 and 10;

50 simulations run for 25 generations (dose 20); 100 simulations run for 20 generations from doses 50 and 100 and 7 simulations run for 25 generations from inoculum dose of 50.

### Supplementary Tables

#### Table S3. Identical isolates found from different geographical groups.

| Carcass ID | Geographical group | Epidemiological cluster (if applicable) |
|---|---|---|
| AN17-01A<br>AN16-165 | Central<br>North | |
| AN16-80<br>LNA2<br>AN16-77 | Central<br>Central<br>North | C1<br>C3 |
| AN16-110<br>AN16-23<br>AN16-122 | Central<br>Central<br>East | |
| AN16-83<br>AN16-109 | Central<br>East | |
| AN16-80<br>AN17-38<br>AN16-160 | Central<br>South<br>South | C1<br><br>C4 |
| AN16-24<br>FL-06<br>FL-MKK | Central<br>South<br>South | |
| AN16-14<br>AN16-23<br>FL16-03 | Central<br>Central<br>South | C2<br>C2 |
| AN16-155<br>AN16-135 | Central<br>South | |

**Table S4. Identical or nearly identical isolates from carcasses sampled several months apart.**

| Groups of related isolates | Isolation dates | SNP differences | Time between sampling |
|---|---|---|---|
| AN16-86-1_T<br>AN16-164_S2 | 29-9-2016<br>25-1-2017 | <br>0 | 4 months |
| AN16-165_S<br>AN17-01-A_B | 26-1-2017<br>30-4-2017 | <br>0 | 3 months |
| AN16-88_D<br>AN16-132_I<br>AN16-132_T | 9-5-2016<br>21-3-2017 | <br><br>0 | 10 months |
| AN16-80_I2/D<br>LNA-2_T1/T2<br>AN16-77_T | 10-5-2016<br>7-10-2016<br>9-10-2016 | <br><br>0 | 5 months |
| AN16-109_D1<br>AN16-83_T1/T2/D<br>AN16-135_S | 18-7-2016<br>20-8-2016<br>20-10-2016 | <br>0/0/1<br>1 | 3 months |
| AN16-80_I1<br>AN16-14S<br>AN16-24_D/T | 10-5-2016<br>16-9-2016<br>26-9-2016 | <br><br>0 | 4 months |
| AN16-110_D3<br>LNA_D3 | 30-6-2016<br>7-10-2016 | <br>0 | 3 months |

SNP: single nucleotide polymorphism

**Table S5. Single nucleotide polymorphism (SNP) distances among pairs of isolates sampled from simulated within-host populations of *Bacillus anthracis*.** Proportion of pairs of isolates in evolved populations with different SNP distances across varying initial inoculum size (dose), sampled in generations 20 and 25, and mean SNP differences across sampled pairs.

| | Generation 20 | | | | | Generation 25[1] | | | | |
|---|---|---|---|---|---|---|---|---|---|---|
| Dose | 0 (%) | 1 (%) | 2 (%) | 3 (%) | Mean SNP distance | 0 (%) | 1 (%) | 2 (%) | 3 (%) | Mean SNP distance |
| 1 | 86.1 | 12.7 | 1.16 | 0.06 | 0.15 | 82.5 | 15.8 | 1.49 | 0.16 | 0.19 |
| 2 | 85.7 | 13.3 | 0.82 | 0.10 | 0.15 | 82.6 | 16.0 | 1.64 | 0.12 | 0.19 |
| 5 | 85.3 | 13.6 | 1.11 | 0.05 | 0.16 | 81.2 | 17.1 | 1.56 | 0.07 | 0.20 |
| 10 | 86.1 | 12.7 | 1.18 | 0.05 | 0.15 | 82.2 | 15.9 | 1.76 | 0.12 | 0.20 |
| 20 | 84.1 | 14.6 | 1.20 | 0.02 | 0.17 | 81.3 | 16.8 | 1.70 | 0.16 | 0.21 |
| 50 | 85.0 | 13.7 | 1.26 | 0.02 | 0.16 | 82.9 | 15.7 | 1.29 | 0.14 | 0.19 |
| 100 | 84.7 | 13.9 | 1.32 | 0.04 | 0.17 | | | | | |

[1] Values for generation 25 with dose 50 are based on a small number of simulations (7) only.

### File S1. Supplementary methods and results.

#### Materials & Methods

##### *Study area*

The population of the Ngorongoro Conservation Area (NCA) is comprised mostly of Maasai pastoralists. The size was based on a population estimate of just over 70,000 inhabitants in 2012 and an annual growth rate of 2.7%. Given this area's conservation status, in addition to livestock, people live in close proximity with various wildlife species.

##### *Research and ethical approval*

Material and data transfer agreements were established as part of the research approvals. This study complied with Tanzania's national access measures for genetic material. Approval and permission to access communities were also obtained from relevant local authorities. Verbal and/or written informed consent was obtained from all owners of livestock sampled, with verbal consent obtained in lieu of written consent where participants preferred. Both verbal and written consent had been approved by the ethical committees.

##### *Sample collection and handling during shipping*

Anthrax was suspected in livestock for any acute mortality where the animal had appeared healthy until the time of death. Terminal bleeding from natural orifices was variably observed. Samples were triple-bagged, with the inner container first disinfected with 10,000 ppm sodium hypochlorite reconstituted from Haz-Tab tablets (Guest Medical, UK). GPS coordinates were taken for samples collected later in the study; for earlier samples, coordinates were estimated by having the field team identify the sampling location on a map of the study area and subsequently plotting these points using Google Earth. When available, GPS coordinates are provided in Table S1 to three decimal places to ensure participant anonymity.

Aliquots from a total of 278 samples from 109 carcasses were shipped to the University of Pretoria, South Africa, for selective culture and DNA extraction from *B. anthracis* isolates. Of the samples sent, about half had previously tested positive by qPCR (142 samples from 63 carcasses). Upon arrival in South Africa, sample tubes were sprayed with disinfectant then heat inactivated for 72 °C for 20 minutes at the Transboundary Animal Diseases section of the Onderstepoort Research Institute as per South African governmental regulations in the control of Foot and Mouth Disease.

***Bacterial isolation***

Tissues

The transit time and heat treatment had resulted in advanced putrefaction and some fungal contamination in the specimen tubes. A salt solution consisting of 900 µL NaCl (5mg/mL) was added to the tubes and left overnight to inhibit the vegetative fungi. The specimens were washed with distilled water and the supernatant discarded four times with successive centrifuging at 8000 XG for 20 min in between. The clean tissue pellet was homogenized in 500 µL phosphate buffered saline (PBS) with glass beads using a single low impact cycle in a Precellys® Tissue homogenizer (Bertin GMBH, Frankfurt Germany).

Swabs

The dry swabs were moistened with 350 µL Dulbecco's saline (fortified with additional 1 M calcium carbonate) for 24 hours. The swabs were softened by the saline solution and the solute took on the appearance of diluted blood after vortexing.

Blood

Blood samples were diluted in 250 µL PBS and were heat treated at 65 °C for 10 minutes to inhibit competition from heat-sensitive bacteria.

Insects

The insects were identified under a stereoscope while being placed in sterile 1.5 mL Eppendorf tubes filled with 350 µL PBS. A homogenizing pestle (Lasec, South Africa) was used to homogenize the sample for plating.

***Sub-Culture and DNA extraction***

Only grey-white, ground-glass textured, non-hemolytic colonies were selected for subculture from sheep blood agar (SBA) (within 24 hours) and dome shaped, rough white colonies were selected from PET (within 48 hours). Differentiating morphology characteristics among suspect isolates included (i) pronounced "medusa head" edges around the colony forming unit; (ii) an uncharacteristic tacky texture when touched with the bacteriologic loop; and (iii) a larger colony diameter than other colony forming units on the same plate.

Once pure, single colonies could be isolated after passage, species was confirmed using a 10 µg penicillin disc (Oxoid) and 10 µL of diagnostic Gamma phage ($5x10^9$ pfu/mL); colonies were identified as *B. anthracis* when demonstrating both penicillin and γ-phage sensitivity after overnight incubation at 37°C, as well as testing positive by qPCR for the protective antigen *pag* gene (BAPA) (3). DNA extraction for qPCR confirmation was performed by harvesting all material on the purity plate in 5 mL of PBS and crude boiled at a 110 °C for 15 minutes. The boiled solution was then centrifuged at 6000 G for 30 min, reserving the supernatant.

To extract genomic DNA for sequencing, a loopful of bacterial colonies from the purity plates was used as the input for the Bioline Isolate II Genomic DNA kit (Meridian Biosciences, UK) with 2 hour 37°C incubation in 20 mg/mL lysozyme (Fluka) solution according to the manufacturer's instructions for Gram-positive bacteria.

***qPCR***

The qPCR for BAPA (3–5) was performed with 2.5 µL DNA in 1x FastStart™ Taq DNA Polymerase mastermix (Roche®) and 0.5 µM of each primer along with 0.2 µM of each FRET probe (Tib MolBiol GmbH) in a final volume of 20 µL. The PCR conditions on a LightCycler™ Nano (Roche®) consisted of an initial cycle at 95 °C for 10 minutes, slope at 20 °C/second, followed by 40 cycles of 95 °C for 10 seconds; 57 °C for 20 second; 72 °C for 30 second, slope 20 °C/second with one single signal acquisition at the end of annealing cycle. Denaturation at 95 °C for 3 seconds with a slope 20 °C/second; 40 °C for 30 seconds, slope 20 °C/second; 80 °C for 3 seconds at a slope of 0.1 °C/second with continuous acquisition of the signal. Cooling to 40 °C for 30 second, slope 20 °C/second (3, 5). Plates and CFU that were negative for both BAPA as well as phage sensitivity were excluded for final nucleic acid extraction selection.

***Additional quality control***

DNA was extracted from 96 isolates and shipped to the UK for further quality control and subsequent sequencing. DNA extracts from 75 isolates from 33 carcasses were submitted for sequencing; these were selected based on having sufficiently high concentration as measured by Qubit (≥1 ng/µl) and lower Ct values (< 25) on qPCR (i.e. higher concentrations of *B. anthracis* DNA); Ct values of DNA extracts from sequenced isolates ranged from 9-21.

***Bioinformatics***

SNP positions identified using VarScan from all individual NCA samples, in addition to A2075, were merged into a single dataset using an in-house python script, resulting in a total of 721 unique variant sites across isolates. Subsequently, read mapping metrics files for the detected set of variant sites were generated for individual isolates using bam-readcount tool v.0.8.0 (https://github.com/genome/bam-readcount). Variant sites were removed based on the following rules: 1) occurred in phage regions as detected by PHASTER (6); 2) occurred in repetitive/homologous genomic regions; 3) more than 3 isolates at a particular site had prevalent base frequency below 89% or/and read depth below 4. In the final alignment file, N character was assigned to sites with prevalent base frequency below 89%, and gap character (-) was assigned to sites having read depth below 4. The resulting filtered alignment file consisted of 611 positions, of which 437 were monomorphic among isolates from the NCA and A2075 (i.e. all isolates had the same allele that differed from the reference). For the purposes of phylogenetic analysis, all gaps and N characters were removed from the alignment file, leaving 499 sites, of which 374 were monomorphic and 125 polymorphic.

***Simulation modelling***

One hundred simulations running for 25 generations were performed for inoculum sizes one, two, five, and ten. Due to computational restrictions, only 50 simulations running for 25 generations were performed for dose 20. One hundred simulations running for 20 generations were performed for doses 50 and 100, while an additional 7 simulations with an initial dose of 50 were run for 25 generations.

### Results

***Bacterial Isolation***

Of the 278 specimens processed for culture, 122 yielded suspect *B. anthracis* colonies, while the remainder had almost no growth on either the *B. anthracis* selective and non-selective media. After testing the suspect colonies with penicillin, Gamma phage and qPCR for the protective antigen *pag* gene, 73 samples (96 individual colonies) had results confirmatory of *B. anthracis* isolates. This included isolates from two carcasses from which none of the samples (soil, tissue and blood) had previously tested positive on qPCR using established Ct value cut-offs (7). After long-term sample storage at ambient temperature, *B. anthracis* isolation was least successful from whole blood, with hardly any bacterial growth after the heat treatment. The tissue, insect, swabs and soil all performed similarly in producing viable isolates, although the swabs had almost pure *B. anthracis* plates for some of the samples, making it the most suitable sample type for bacterial isolation under this type of collection and storage conditions. The 122 specimens that yielded any growth at all were almost exclusively made up of a combination of *B. endophyticus, B. cereus, B. pumilus, B. megaterium, B. subtilis, B. thuringiensis* and *B. anthracis* representing the hardiest spore formers within the specimens.

Several *B. anthracis* isolates were obtained from flies captured on or around carcasses, most of which were *Muscidae* species. While speciation was difficult in some cases due to damaged wings and/or proboscis, species were predominantly *Musca crassirostris* and to a lesser extent *Musca domestica.* No efforts were made to determine whether the spores were on the surface or in the gut of the flies. In certain ecosystems, flies are thought to play an important role as mechanical vectors, spreading spores onto leaves which may become a source of infection for browsing animals (8, 9). The role of biting flies as a source of *B. anthracis* infection is less well defined (3).

***Sequence quality***

GC content of the raw reads was unexpectedly low (<34; n=1) or high (>36; n=5), and/or total *de novo* assembly length was above the expected length of ~5.5 MB (n=13), and/or there were a high number of contigs (>1000; n=11), indicative of challenges with assembly that could be related to mixed culture. A total of 16 isolates had one or more of these issues (S1 Table). Strict filtering criteria were implemented during reference-based mapping to address this issue.

***Simulation modelling***

After 20 generations and across 100 simulations, a population with a starting dose of 1 (n = 524,288) had an average of 2,258 (range 2,140 to 2,364) unique SNP profiles, i.e. that differed by at least one SNP. This rose to 71,845 (range 71,049 to 72,610) after 25 generations. When averaging across simulations, inoculum size (infectious dose) did not significantly influence the proportion of the population identical to the infecting genome, or that differed by 1 or 2 SNPs. Across inoculum sizes, the proportion of genomes identical to the infecting dose in our simulations tended to remain high, and while genomes differing by a maximum of 5 (doses 1 and 2) and 6 (doses 5 – 100) SNPs were observed, these were incredibly rare and therefore unlikely to be sampled; the majority of variant genomes differed from the inoculum genome by a single SNP. Greater variability around the average proportion of the population with different numbers of SNPs was observed in populations simulated from the smaller inoculum sizes due to increased impact of stochasticity in early generations. For example, in a single simulation with an inoculum size of 1, a mutation emerged in the first replication and therefore no genomes identical to the infecting dose were observed in further generations (Fig. S8-A), while in some simulations initiated with an infectious dose of 2, two major variants separated by a single SNP were both present at fairly even proportions (Fig. S9-B).

Proportions of sampled pairs of genomes from the simulated populations with different numbers of SNP differences are summarized in Table S5, which also shows the mean SNP distance averaged across sampled pairs. This mean SNP distance ranged from 0.15 (dose = 1) to 0.17 (dose = 100). While pairwise distances ranging from 0 to 4 were observed, this only included 33 observations of 3 SNP distance (0.05%) and 5 observations of 4 SNP distance (0.008%).

The relationship between higher dose and higher mean SNP distance was statistically significant ($p < 0.005$) though the estimated effect size was small (+0.0014 per extra 10 genomes in dose) and resulted in only a slight increase in the chance of drawing pairs of genotypes with greater SNP distances. For example, the proportion of samples with SNP distances of 2 or more increased only from 1.22% with an inoculum size of 1, to 1.37% with an inoculum of 100 genomes.

At least 50 simulations initiated with inoculums of 1, 2, 5 10 and 20 bacteria were run for an additional 5 generations. The mean SNP distance between pairs sampled in generations 20 to 25 tended to increase slightly (Fig. 6D), by around 0.0085 per generation ($p < 0.0001$). This gradual increase resulted in the proportion of samples with a SNP distance of 2 rising from 1.15% in generation 20 to 1.74% in generation 25. In 270,000 pairs sampled in generations 21 to 25, a SNP distance of 4 was observed only 8 times and a SNP distance of 5 was sampled once (in a population simulated from an infectious dose of 10 sampled in the 24$^{th}$ generation). Such an observation is incredibly rare as it requires, in the most likely scenario, sampling 2 genotypes differing from the inoculum by 2 and 3 non-shared mutations.
